# Impaired SorLA maturation and trafficking as a new mechanism for *SORL1* missense variants in Alzheimer disease

**DOI:** 10.1101/2021.06.18.448296

**Authors:** Anne Rovelet-Lecrux, Sebastien Feuillette, Laetitia Miguel, Catherine Schramm, Ségolène Pernet, Isabelle Ségalas-Milazzo, Laure Guilhaudis, Stéphane Rousseau, Gaëtan Riou, Thierry Frébourg, Dominique Campion, Gaël Nicolas, Magalie Lecourtois

**Affiliations:** 1Normandie Univ, UNIROUEN, Inserm U1245, CHU Rouen, Department of Genetics and CNR-MAJ, FHU G4 Génomique, F-76000 Rouen, France; Normandie Université, UNIROUEN, INSA de Rouen, CNRS, Laboratoire COBRA (UMR 6014 & FR 3038), Rouen – France; Inserm U1234, Flow Cytometry Core – IRIB, Rouen, France

**Keywords:** SorLA, iPSC, Alzheimer’s disease, maturation defects, trafficking

## Abstract

The SorLA protein, encoded by the *SORL1* gene, is a major player in Alzheimer’s disease (AD) pathophysiology. Functional and genetic studies demonstrated that SorLA deficiency results in increased production of Aβ peptides, and thus a higher risk of AD. A large number of *SORL1* missense variants have been identified in AD patients, but their functional consequences remain largely undefined. Here, we identified a new pathophysiological mechanism, by which rare *SORL1* missense variants identified in AD patients result in altered maturation and trafficking of the SorLA protein. An initial screening, based on the overexpression of 71 SorLA variants in HEK293 cells, revealed that 15 of them (S114R, R332W, G543E, S564G, S577P, R654W, R729W, D806N, Y934C, D1535N, D1545E, P1654L, Y1816C, W1862C, P1914S) induced a maturation and trafficking-deficient phenotype. Three of these variations (R332W, S577P, and R654W) and two maturation-competent variations (S124R and N371T) were further studied in details in CRISPR/Cas9-modified hiPSCs. When expressed at endogenous levels, the R332W, S577P, and R654W SorLA variants also showed a maturation defective profile. We further demonstrated that these variants were largely retained in the endoplasmic reticulum, resulting in a reduction in the delivery of SorLA mature protein to the plasma membrane and to the endosomal system. Importantly, expression of the R332W and R654W variants in hiPSCs was associated with a clear increase of Aβ secretion, demonstrating a loss-of-function effect of these SorLA variants regarding this ultimate readout, and a direct link with AD pathophysiology. Furthermore, structural analysis of the impact of missense variations on SorLA protein indicated that impaired cellular trafficking of SorLA protein could be due to subtle variations of the protein 3D structure resulting from changes in the interatomic interactions.

## INTRODUCTION

Alzheimer’s disease (AD) is a genetically heterogenous disorder. Besides rare families with autosomal dominant early-onset AD (EOAD, onset before 66 years) due to pathogenic variants in *APP*, *PSEN1* or *PSEN2*, the determinism of AD is complex and includes a large multigenic component. A wide diversity of genetic risk factors has been identified, varying both in terms of frequencies and strength of effect on AD risk (Nicolas *et al*, 2016a). Exome and genome sequencing recently allowed the identification of three genes harboring a higher burden of rare non-synonymous variants in AD cases as compared to controls: *TREM2*, *SORL1*, and *ABCA7*.

The *SORL1* gene (Sortilin-related receptor 1) encodes a ∼250-kDa transmembrane protein, termed Sortilin-related receptor (SorLA), with multiple functional domains. The ectodomain of SorLA is a complex mosaic structure comprising a VPS10p domain, a Sortilin C domain, LDLR class B and class A repeats, an EGF-like domain, and a cassette of six fibronectin type-3 domains (Figure 1A). Newly synthesized SorLA molecules follow the constitutive secretory pathway from the endoplasmic reticulum (ER) to the cell surface, through the Golgi. During their transport through the secretory pathway, SorLA proteins are subjected to several post-translational modifications, including both N- and O-glycosylation, and maturation processes (Jacobsen *et al*, 2001; Fiete *et al*, 2007; Schmidt *et al*, 2007; Fjorback *et al*, 2012; Steentoft *et al*, 2013; Hampe *et al*, 2000; Christensen *et al*, 2020), resulting in the presence of immature and mature species. At the cell surface, a minority of SorLA molecules are subject to proteolytic shedding, releasing the soluble ectodomain of the receptor, termed soluble SorLA (sSorLA). However, most SorLA molecules remain intact and undergo clathrin-dependent endocytosis. Internalized molecules move to the early endosomes from where most of them will sort to the trans-Golgi network (TGN) to continuously shuttle between TGN and endosomes thereafter.

**Figure 1:**
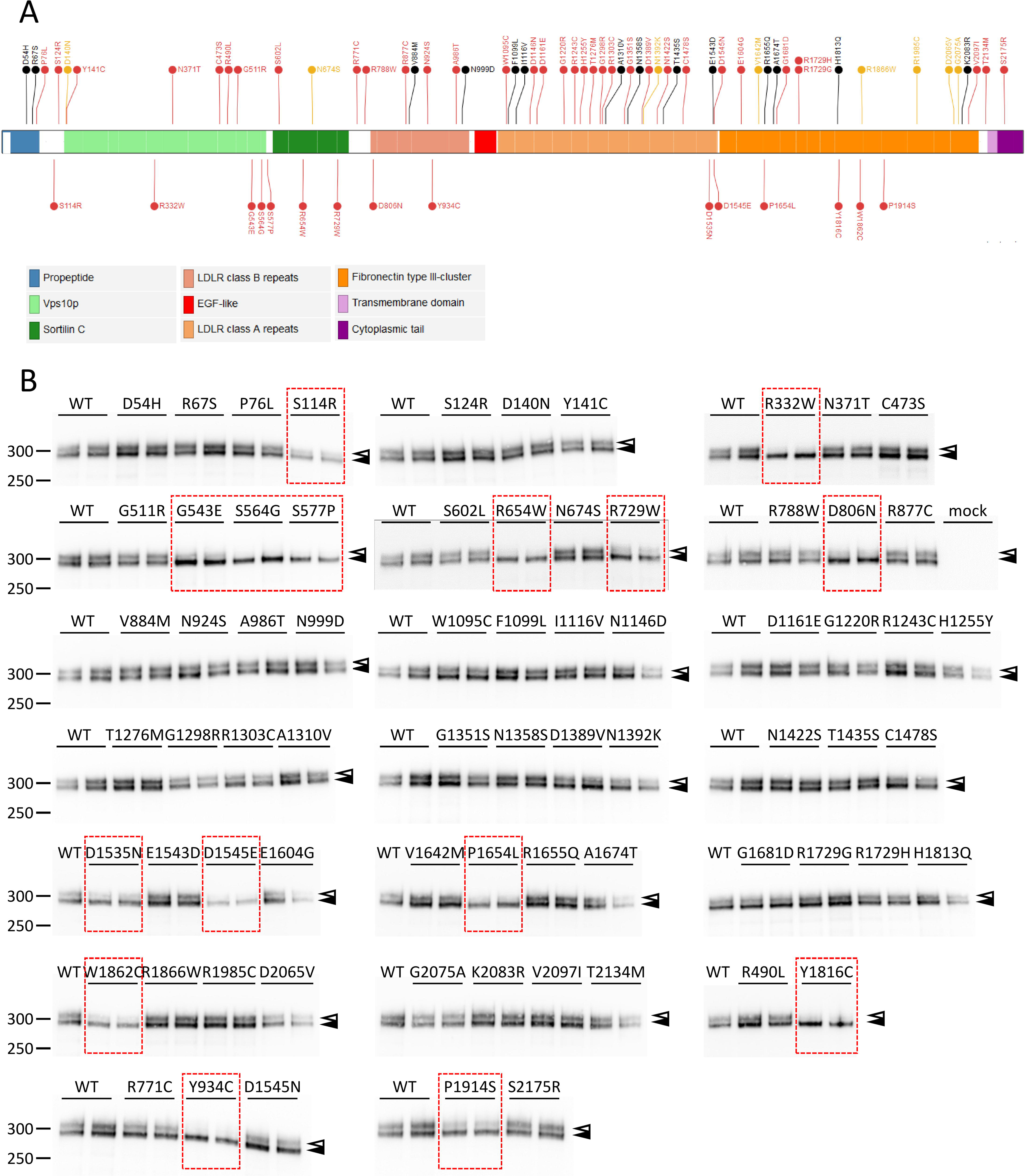
Identification of SorLA missense variants that reduce production of the complex-glycosylated mature form of the protein in HEK293 cells. (A) For the schematic presentation of SorLA domains structure, we used the domains provided by Uniprot (https://www.uniprot.org/uniprot/Q92673) and added the Sortilin C domain provided by Pfam (https://pfam.xfam.org/protein/Q92673): signal peptide (a.a. 1-28), propeptide (a.a. 29-81), Vps10p (BNR repeats) (a.a. 136-573), Sortilin C (a.a. 588-752), LDLR class B repeats (a.a. 800-1013), EGF-like (a.a. 1026-1072), LDLR class A repeats (a.a. 1076-1551), Fibronectin type III-cluster (a.a. 1557-2118), Transmembrane domain (a.a. 2138-2158) and Cytoplasmic tail (a.a. 2159-2214). The Mis3, Mis2, and Mis0-1 missense variants examined in this study are shown in red, orange and black, respectively. The 15 Mis3 variants presenting maturation defects are placed below the schematic presentation of the SorLA protein. (B) Western Blot analyses of the SorLA protein maturation profile in HEK293 cells transiently expressing wild-type (WT) and missense variants of SorLA. For each variation, the pattern was confirmed in at least 2 independent replicates. Blots were probed with an anti-SorLA antibody and representative blots are presented. Immature core-glycosylated and mature complex-glycosylated SorLA are indicated with solid and empty arrowheads, respectively. The molecular masses of marker proteins in kDa are shown on the left. Maturation-deficient SorLA variants are highlighted with a red rectangle.

Several studies have highlighted a protective function of wild-type SorLA against AD. Indeed, loss of *SORL1* expression in several mouse models of AD increases extracellular Aβ levels, the aggregation of which triggers AD pathology (Andersen *et al*, 2005; Dodson *et al*, 2008; Rohe *et al*, 2008). On the other hand, overexpression of *SORL1* in neuronal and non-neuronal cell lines limits the processing of APP, the Aβ precursor protein, and decreases Aβ production (Andersen *et al*, 2005; Offe *et al*, 2006; Rogaeva *et al*, 2007; Schmidt *et al*, 2007). The known mechanisms whereby SorLA acts to lower Aβ production have been reviewed recently in details (Andersen *et al*, 2016; Schmidt *et al*, 2017). First, SorLA is involved in the trafficking and recycling of APP. By binding to the retromer complex, SorLA redirects APP from the early endosomes back to the Golgi apparatus through the recycling pathway. This in turn results to reduced amyloidogenic processing of APP, by preventing it from being targeted to the late endosome compartments, where APP is cleaved into Aβ by the β- and γ-secretases. Second, SorLA may directly bind nascent Aβ peptides and target them to lysosomes, which also contributes to the reduction of Aβ secretion (Caglayan *et al*, 2014).

Rare coding *SORL1* variants have been found in ∼3% of AD patients and are now considered a major AD risk factor (Nicolas *et al*, 2016b; Bellenguez *et al*, 2017; Pottier *et al*, 2012; Campion *et al*, 2019). The association has been demonstrated at the gene level by aggregating protein-truncating (PTV, all of which are ultra-rare) and rare (allele frequency below 1%), predicted damaging, missense *SORL1* variants. The interpretation of the effect of PTV appears quite straightforward, i.e. leading to reduced SorLA levels through nonsense-mediated decay (NMD) and/or production of truncated proteins and hence increased Aβ production. However, the functional effect of missense variants and their pathophysiological mechanisms are hard to predict. Whether or not a given missense variant behaves as a loss-of-function variant and by which mechanisms remain to be determined for most of them. At the post-genomic era, interpretation of missense variants remains a major obstacle for many genes, including *SORL1*. Hence, most *SORL1* missense variants can basically not be used in a clinical setting, up to now. To date, only 6 missense variants have been studied *in vitro* and have shown varying degrees of decrease in SorLA protein function (Caglayan *et al*, 2014; Vardarajan *et al*, 2015; Cuccaro *et al*, 2016). Among them, demonstration of an associated mechanism was shown for only one, p.Gly511Arg, which diminished the ability of mutant SorLA to target nascent Aβ peptides to lysosomes. Other missense variants may act by reducing SorLA binding to other partners, including APP, or by resulting in an unstable, misfolded or immature protein.

Here, we analyzed 71 rare missense variants identified from exome sequencing in AD patients by our group. An initial screening, based on overexpression in HEK293 cells, revealed that 15 of them showed a maturation and trafficking-deficient phenotype. Three of these variations and two maturation-competent variations were further studied in details on endogenous *SORL1* in CRISPR/Cas9-modified hiPSCs, validating the results obtained in HEK293 cells and highlighting gradual effects of the variations. We also demonstrated that two variants affecting the proper maturation of SorLA were associated with an increase of Aβ secretion in the cellular medium, thus providing a direct link with AD pathophysiology. Finally, structural analysis revealed that this impaired cellular trafficking could result from subtle variations of the protein 3D structure and changes in the interatomic interactions.

## RESULTS

### Identification of SorLA variants that reduce the level of complex-glycosylated mature SorLA protein in HEK293 cells

In our group, exome sequencing of 1383 patients with AD (97% EOAD) led to the identification of 83 missense *SORL1* variants (Bellenguez *et al*, 2017; Campion *et al*, 2019; Holstege *et al*, 2020). We studied 71 of these missense variants, distributed all along the SorLA protein (Figure 1A) and classified according to their predicted deleteriousness assessed by three bioinformatic tools: PolyPhen-2, MutationTaster and SIFT. Forty-nine of them were predicted damaging by all three bioinformatics tools (Mis3 variants), 8 were predicted damaging by two of the three tools (Mis2) and 14 were predicted damaging by one or none of the tools (Mis1-0) (Figure 1A).

To study the effect of these rare *SORL1* missense mutations on SorLA protein maturation, cDNA constructs encoding the full-length wild-type (WT)-SorLA or SorLA variants identified in AD patients were used to transfect HEK293 cells for transient expression. When overexpressed in HEK293 cells, WT-SorLA protein consisted of two distinct forms on immunoblot analysis (Figure 1B, Figure 2) that were hypothesized to be different glycoforms of the protein. As previously shown (Jacobsen *et al*, 2001; Christensen *et al*, 2020), removal of all *N*-linked glycans by treatment with PNGase F resulted in a single band that migrates at ∼250kDa, which corresponds to the predicted molecular weight of the full-length SorLA (SorLA^FL^) protein (Figure 2A). Treatment with neuraminidase altered the mobility of only the upper band that subsequently migrated as the lower form, indicating that the upper species correspond to complex type *N*-glycans capped with sialic acid residues (Figure 2A). No further shift in the migration was seen while combining neuraminidase and O-glycosidase (Figure 2A), or O-glycosidase and PNGase F (Figure 2B), suggesting that SorLA lacks O-glycosylation in HEK293 cells. The conversion of high-mannose type *N*-glycans to complex *N*-glycans occurs in the medial Golgi region. When proteins are correctly processed through the endoplasmic reticulum (ER) and Golgi, *N*-glycans become resistant to Endo H. Sensitivity to Endo H therefore indicates the presence of proteins bearing high mannose *N*-glycans that still reside in the ER. In our experiments, Endo H digestion yielded two bands with reduced molecular size (Figure 2C). As expected, the upper band that corresponds to mature complex type *N*-glycans capped with sialic acid residues, was somewhat resistant to Endo H. However, the lower SorLA species were equally sensitive to treatment with Endo H and PNGase F yielding a band at ∼250kDa, indicating that they correspond to species bearing high mannose *N*-glycans. Together, these data indicated that SorLA exist as two species in HEK293 cells: a mature complex-glycosylated form of SorLA that corresponds to the upper band of the doublet and an immature oligomannoside *N*-linked core-glycosylated form of the protein that corresponds to the lower band.

**Figure 2:**
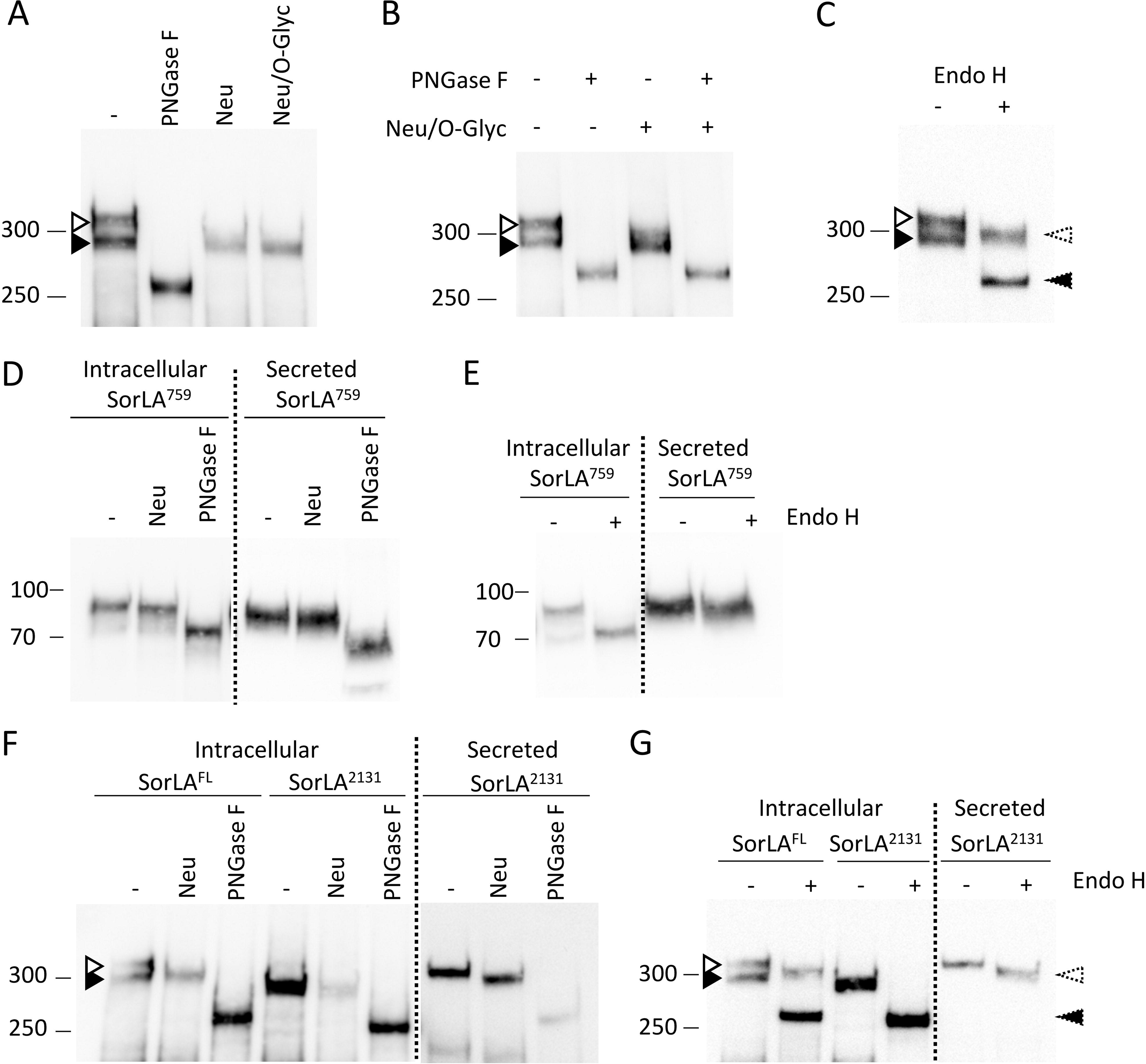
Characterization of the maturation process of SorLA proteins in HEK293 cells. Western blot analyses after glycosidase treatments of wild-type full-length (SorLA^FL^) (A, B) or secreted truncated (SorLA^759^ or SorLA^2131^) (C-G) forms of SorLA proteins transiently overexpressed in HEK293 cells. Blots were probed with an anti-SorLA antibody and representative blots are presented. Cellular lysates (intracellular) or supernatants (secreted) were treated in the absence or presence of the glycosidases PNGase F, Neuraminidase (Neu), O-Gycosidase (O-Glyc). Untreated samples (–) were incubated under the same conditions but without enzymes. PNGase F cleaves both high-mannose, hybrid and complex *N*-glycans from glycoproteins. Endo H cleaves preferentially high-mannose type *N*-glycans, but not complex *N*-glycans. Neuraminidase cleaves sialic acid from either complex or O-glycans. O-glycosidase removes O-linked glycans, but this cleavage is more effective only after removing terminal sialic residues by a neuraminidase. The positions of immature core-glycosylated and mature complex-glycosylated forms of SorLA are indicated with solid and empty arrowheads, respectively. Resulting corresponding species obtained after neuraminidase treatment were indicated with dashed arrowheads. The molecular masses of marker proteins in kDa are shown on the left.

Of the 71 missense variants studied, 56 displayed a maturation profile similar to the one observed for WT-SorLA (Figure 1B). In contrast, for 15/71 (15/49 of Mis3 variants (∼30%)), the upper band corresponding to the glycoform of the protein bearing complex-type *N*-glycans was fainter or barely detectable (S114R, R332W, G543E, S564G, S577P, R654W, R729W, D806N, Y934C, D1535N, D1545E, P1654L, Y1816C, W1862C, and P1914S), indicating defects in the maturation of complex *N*-glycans for these variants. These data are summarized in Table S1.

### SorLA variations associated with abnormal immunoblot profiles interfere with SorLA trafficking to the plasma membrane in HEK293 cells

The reduced level of fully mature SorLA could be associated with trafficking defects of the protein along the secretory pathway, thereby preventing the trafficking of SorLA to the cell surface. To explore the impact of *SORL1* variations on SorLA trafficking towards cytoplasmic membrane, we took advantage of constructs leading to secreted truncated forms of SorLA, comprising different parts of the luminal part of SorLA, but not its cytoplasmic tail or transmembrane segment. These constructs encoded either amino acids 1-759 (SorLA^759^) or amino acids 1-2131 (SorLA^2131^) of the SorLA protein (Figure 3A) (Jacobsen *et al*, 2001). After transient transfection in HEK293 cells, intracellular or secreted WT-SorLA proteins were subjected to Western blotting. The WT-SorLA^759^ and WT-SorLA^2131^ proteins were detected both in the cellular lysate and in the medium (Figure 3B, Figure S1, Figure S2), indicating that both truncated constructs reached the plasma membrane and were efficiently secreted. Treatment of WT SorLA^2131^ and SorLA^759^ proteins with glycosidase showed that the species present in the lysate were immature high-mannose type *N*-glycans (PNGase F and Endo H sensitive, Neu insensitive), whereas the secreted forms of the protein corresponded to mature complex *N*-linked glycoproteins (PNGase F and Neu sensitive, Endo H partially resistant) (Figure 2D-G).

**Figure 3:**
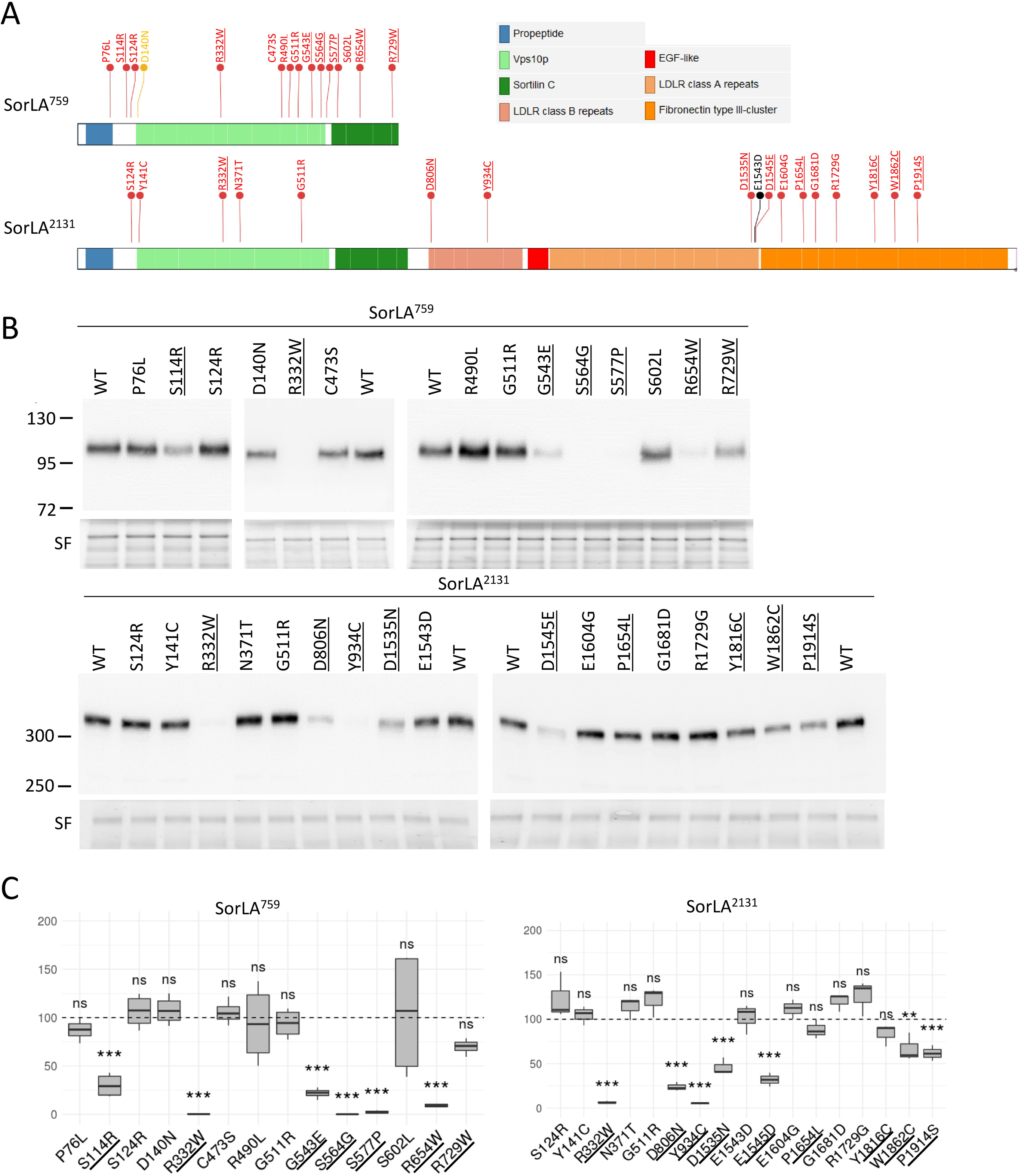
SorLA variations associated with abnormal immunoblot profiles interfere with the secretion of SorLA truncated constructs in HEK293 cells. (A) Schematic representation of the soluble truncated SorLA^759^ and SorLA^2131^ proteins, and the location of SorLA variations examined. The Mis3, Mis2, and Mis0-1 missense variants studied are shown in red, orange and black, respectively. Maturation-deficient SorLA variants are underlined. (B) Wild-type (WT) and mutants SorLA proteins were expressed in HEK293 cells. The SorLA proteins secreted into the cellular medium were analyzed by immunoblotting using an anti-SorLA antibody. Total protein was used as the loading control with Stain-free (SF) technology. Representative blots are presented. Maturation-deficient SorLA variants are underlined. The molecular masses of marker proteins in kDa are shown on the left. (C) Quantification of secretion efficiency from 4 independent experiments (SorLA^759^) or 3 independent experiments (SorLA^2131^). The normalized quantification of the SorLA proteins in the cellular medium is reported in the graphs. For both SorLA^759^ and SorLA^2131^, WT-SorLA were arbitrarily set at 100 arbitrary units. Protein levels were compared by using a regression model adjusted for experiment as random effect. P-value significances are displayed after Bonferroni’s correction for 14 and 17 statistical tests for SorLA^759^ and SorLA^2131^, respectively.

The 15 missense variants corresponding to maturation-defective proteins in the context of the full-length protein (S114R, R332W, G543E, S564G, S577P, R654W, R729W, D806N, Y934C, D1535N, D1545E, P1654L, Y1816C, W1862C, and P1914S) and 13 missense variants presenting a wild-type maturation profile (P76L, S124R, D140N, Y141C, N371T, C473S, R490L, G511R, S602L, E1543D, E1604G, G1681D, and R1729G) were introduced into the WT-SorLA^759^ and WT-SorLA^2131^ constructs by site-directed mutagenesis (Figure 3A). All assessed SorLA variants were detected in the cellular lysate, indicating that they were properly expressed in HEK293 cells (Figure S1). The truncated constructs carrying the S114R, R332W, G543E, S564G, S577P, R654W, D806N, Y934C, D1535N, D1545E, W1862C or P1914S variant, associated with a deficient maturation profile using the full-length construct, all showed a statistically significant lower level of secretion compared to their corresponding wild-type construct (Figure 3B, 3C, Table S1, S5). The R729W variant also resulted in a lower level of secretion, but did not reach statistical significance. The P1654L and Y1816C variants did not show statistically significant difference compared to the WT-SorLA^2131^. Of the 13 missense variants that displayed a wild-type maturation profile (P76L, S124R, D140N, C473S, R490L, G511R, S602L, E1543D, E1604G, G1681D, R1729G), all presented a level of secretion similar to the wild-type form of their matching constructs. Note that some variations (S124R, R332W, G511R) have been introduced both in the SorLA^759^ and SorLA^2131^ constructs and yielded similar quantitative results. The results obtained using secreted truncated forms of SorLA suggest that maturation defects of glycosylation are associated with a default of protein trafficking towards the plasma membrane.

To validate these results in the context of the full-length SorLA protein, we evaluated the amount of SorLA at the plasma membrane by cell-surface biotinylation experiments (Figure 4). Compared to the wild-type form of the protein, all SorLA variants exhibiting maturation defects (S114R, R332W, G543E, S564G, S577P, R654W, R729W, D806N, Y934C, D1535N, D1545E, P1654L, Y1816C, W1862C, P1914S) showed a lower amount of biotinylatable cell-surface SorLA, with a less drastic effect for the variants located in the Fibronectin-type III domain. The cytosolic FUS protein was used as a control to exclude non-specific labelling of the intracellular fractions. As expected, no FUS staining was detected in the biotinylated fraction.

**Figure 4:**
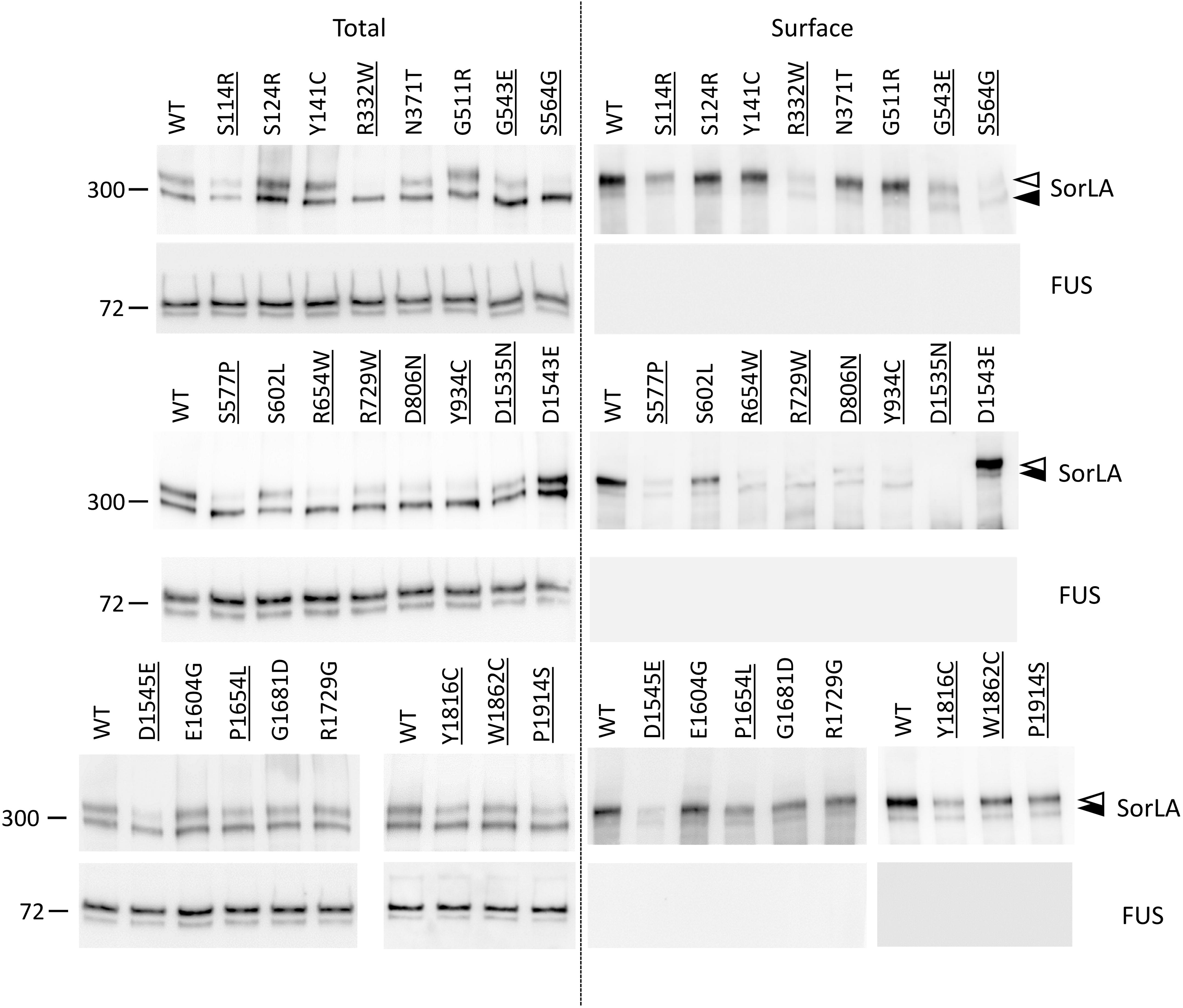
Maturation defective SorLA variants display reduced cell-surface levels of SorLA protein in HEK293 cells. Surface biotinylation experiments to examine cell-surface steady-state levels of SorLA protein in HEK293 cells expressing wild-type (WT) and SorLA variants. Total lysates (Total) and biotinylated fraction**s** (Surface) were analyzed by immunoblotting using an anti-SorLA antibody. FUS, an intracellular protein, was used as an internal control. The absence of FUS staining in the biotinylated fraction demonstrates the integrity of the plasma membrane during the biotinylation experiment. The molecular masses of marker proteins in kDa are shown on the left. Maturation-deficient SorLA variants are underlined. The positions of immature core-glycosylated and mature complex-glycosylated SorLA^FL^ are indicated with solid and empty arrowheads, respectively.

Altogether, these data show that maturation defects are associated with an impairment of SorLA trafficking to the cell surface.

### SorLA variations interfere with SorLA maturation and trafficking to the plasma membrane in hiPSC

To validate the functional effect of rare missense variations in the context of endogenous *SORL1* expression, we used CRISPR/Cas9 gene editing to introduce three maturation-defective variants (R332W, S577P and R654W) and two variants showing a wild-type maturation profile (S124R and N371T) into a control hiPSC line previously generated by our group. Sanger sequencing of colonies derived from single putatively edited cells confirmed the presence of the S124R, R332W, N371T, S577P and R654W variants at the homozygous state in each respective cell line (Figure S2).

We first assessed SorLA maturation profile in CRISPR/Cas9-edited hiPSC and isogenic wild-type control cells by Western blot analyses. In the wild-type cell lines, the SorLA protein displayed a two-band migration profile, with the upper band being the overwhelming majority (Figure 5A). Glycosidase treatment confirmed that the upper band corresponded to mature complex N-linked glycoproteins (PNGase F and Neu sensitive, Endo H partially resistant), and the lower band to immature high-mannose type *N*-glycans (PNGase F and Endo H sensitive, Neu insensitive) (Figure 5A). The S124R and N371T hiPSC lines showed a SorLA migration profile similar to that observed for the wild-type cell lines (Figure 5B). On the other hand, in the R654W hiPSC lines, we observed a drastic downwards mobility shift of the SorLA proteins. Regarding the R332W and the S577P clones, we detected an intermediate profile, with a decrease in the intensity of the upper band and concomitantly an increase in the intensity of the lower band, appearing to be more pronounced for the R332W line (Figure 5B). These patterns were confirmed in three independent hiPSC clones, indicating that the results were not clone-dependent.

**Figure 5:**
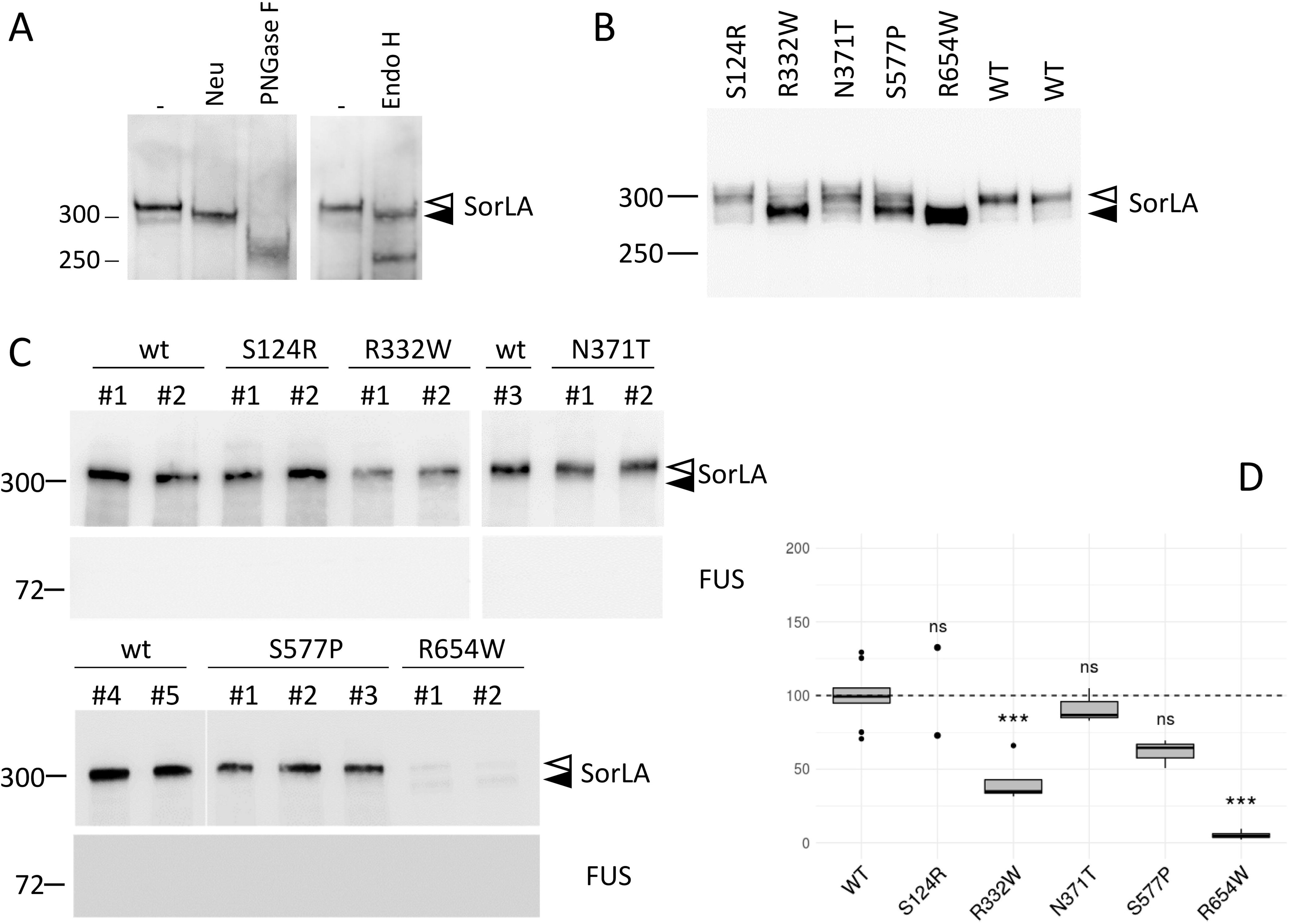
Missense variants alter the maturation and cell-surface levels of SorLA protein in hiPSCs. (A) Western blot analyses of glycosidase treatments of wild-type full-length (SorLA^FL^) expressed in hiPSC at endogenous level. Cellular lysates were treated in the absence or presence of the glycosidases PNGase F, Neuraminidase (Neu), O-Gycosidase (O-Glyc). Untreated samples (–) were incubated under the same conditions but without enzymes. (B) Maturation profile of wild-type and SorLA missense variants expressed in hiPSC at endogenous level. (C) Surface biotinylation experiments to examine cell-surface steady-state level of SorLA protein in wild-type or S124R, R332W, N371T, S577P and R654W CRISPR/Cas9-edited hiPSC. The cytosolic FUS protein was used as control to exclude non-specific labelling of the intracellular fractions. Several independent clones (#) are presented. (A-C) Blots were probed with an anti-SorLA antibody and representative blots are presented. Immature core-glycosylated and mature complex-glycosylated SorLA are indicated with solid and empty arrowheads, respectively. The molecular masses of marker proteins in kDa are shown on the left. (D) Normalized quantification from at least 2 independent experiments. WT mean was arbitrarily set at 100 units. Protein levels were compared by using a regression model adjusted on the experiment as random effect. P-value significances are displayed after Bonferroni’s correction for 5 statistical tests.

In HEK293 cells, altered maturation profile correlated to defects in SorLA trafficking to the plasma membrane, and lower SorLA levels at the plasma membrane. We thus performed cell-surface biotinylation experiments on wild-type and CRISPR/Cas9-modified hiPSCs. In wild-type cells, only the mature band was present at the plasma membrane (Figure 5C, Figure S3), suggesting that SorLA maturation is a prerequisite to its proper trafficking to the plasma membrane. The S124R and N371T variants presented a level of plasma membrane-associated SorLA similar to the wild-type protein (Figure 5, C, D, Table S5). In contrast, 2 of the 3 maturation-defective variants (R332W and R654W) showed statistically significant lower levels of mature SorLA protein at the plasma membrane. The presence of the S577P variant also resulted in lower levels of SorLA at the plasma membrane, but levels were not statistically different from the wild-type form after correction for multiple testing. The amount of SorLA protein at the plasma membrane correlated with the amount of the mature form, the R654W showing the strongest effect with almost no detectable protein at the plasma membrane. The cytosolic FUS protein was used as control to exclude nonspecific labelling of the intracellular fractions. As expected, no FUS staining was detected in the biotinylated fraction.

Altogether, these data obtained in hiPSC lines expressing the R332W, S577P and R654W missense *SORL1* variants at endogenous levels confirmed that these SorLA variants exhibit diverse degrees of maturation defects and lower levels of mature protein at the plasma membrane.

### Retention of the SorLA maturation defective variants in endoplasmic reticulum

As SorLA maturation occurs during its trafficking from the endoplasmic reticulum towards the plasma membrane through the Golgi apparatus, we hypothesized that maturation-defective SorLA variants should present a disturbed subcellular localization with a redistribution within organelles involved in the secretory pathway. To address this issue, we performed immunofluorescence experiments and colocalization analyses on WT or CRISPR/Cas9-edited hiPSCs.

In accordance with previous reports (Madsen *et al*, 2019; Nielsen *et al*, 2007; Gustafsen *et al*, 2013), SorLA immunostaining in WT hiPSCs revealed a polarized localization, both in large perinuclear vesicles, and in smaller vesicles dispersed throughout the cytoplasm (Figure 6). In the S124R and N371T hiPSCs, the subcellular distribution of SorLA protein was similar to that observed in WT cells. In contrast, the R332W and R654W hiPSCs presented only small SorLA-positive vesicles with widespread distribution throughout the cytoplasm. The S577P hiPSCs showed an intermediate distribution of SorLA, with fewer large vesicles close to the nucleus and larger amounts of small vesicles dispersed in the cytoplasm compared to the WT cells. *SORL1* knockout (KO) hiPSCs were used as a negative control (Figure 6, Figure S5).

**Figure 6:**
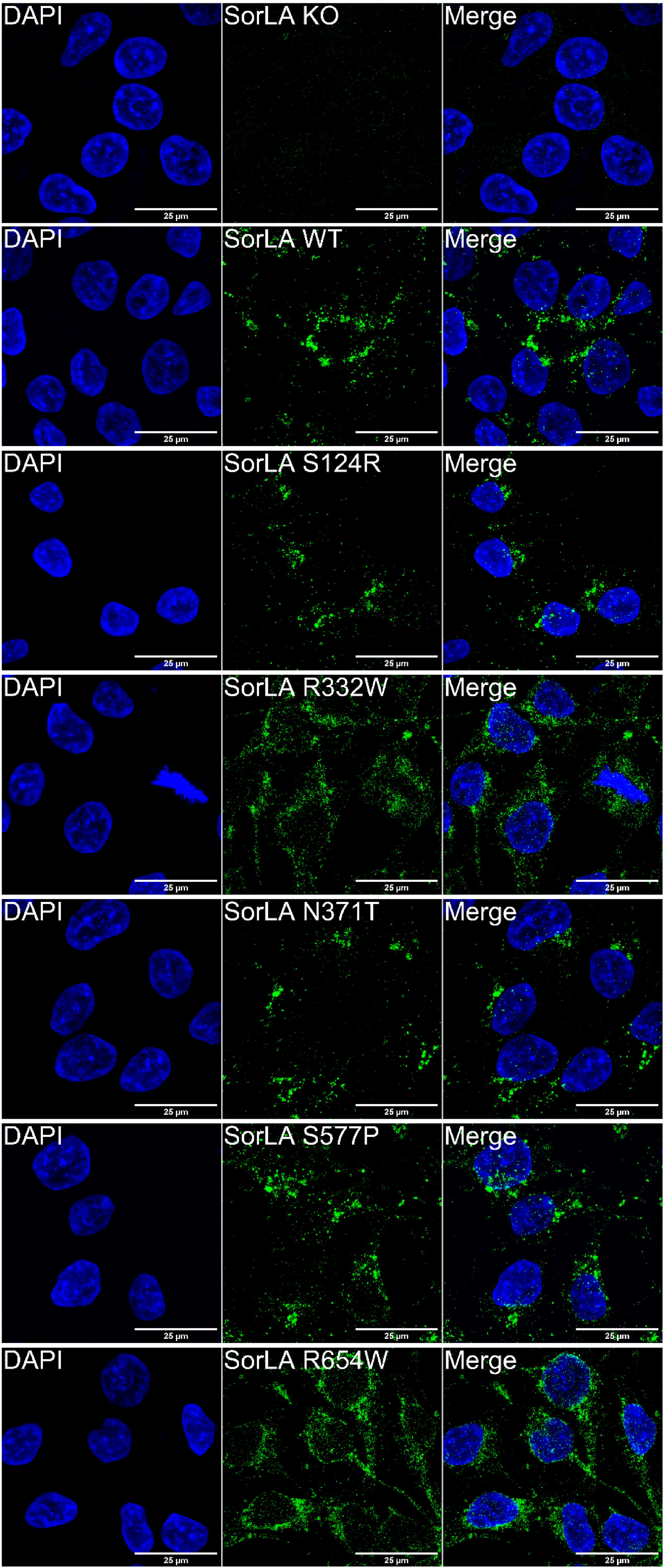
Subcellular localization of the SorLA protein in wild-type and CRISPR/Cas9-edited human iPSC lines. In green, SorLA immunostaining. In blue, DAPI counter-staining.

Next, we performed immunofluorescence co-labelling of SorLA with Cyclophilin B and Rab5 as markers of the endoplasmic reticulum and the early endosomes, respectively (Figure S5, Figure S6). The extent of SorLA colocalization with the two markers was assessed by calculating the M1 Manders’ coefficient (Bolte & Cordelières, 2006) (Figure 7, Figure S7). Regarding the overlap of SorLA with the Cyclophilin B, the M1 coefficients did not statistically differ between the WT, S124R and N371T hiPSCs, indicating similar proportions of SorLA in the endoplasmic reticulum. In contrast, we observed statistically higher M1 coefficients for the R332W, R654W and S577P hiPSCs, indicating a higher proportion of SorLA proteins in the endoplasmic reticulum. Note that the variation of the M1 coefficient was less pronounced for the S577P variant (Figure 7A). Regarding the overlap of SorLA with Rab5, we observed inverse variations of the M1 coefficients. Accordingly, the M1 coefficients did not statistically differ between the WT, S124R and N371T hiPSCs, whereas the M1 coefficients were statistically smaller in the R332W, R654W and S577P hiPSCs compared to the WT cells, indicating a lower proportion of SorLA in early endosomes. Again, the variation of the M1 coefficient was less pronounced for the S577P variant (Figure 7B).

**Figure 7:**
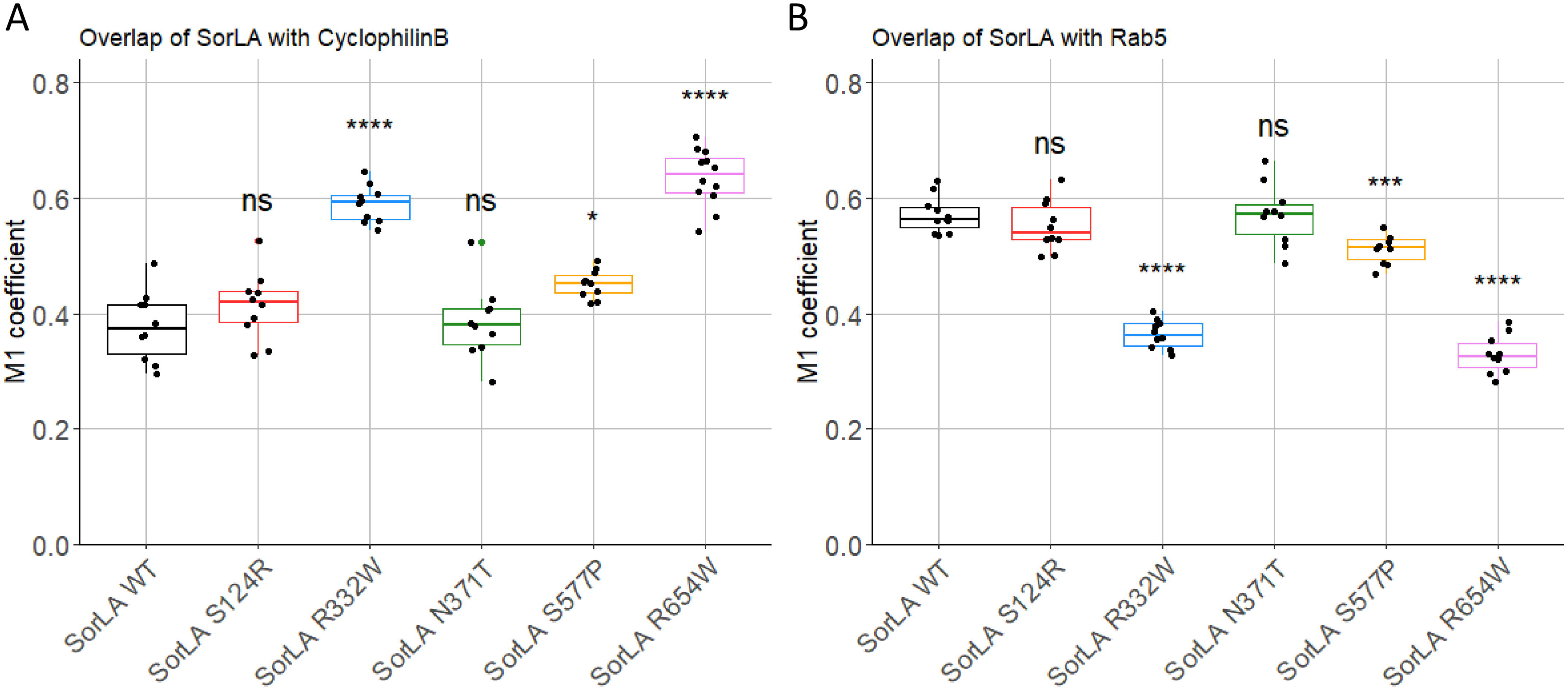
Colocalization analyses of SorLA with markers of the endoplasmic reticulum and early endosomes in wild-type and CRISPR/Cas9-edited human iPSC lines. Overlap of SorLA staining with cyclophilin B (A) and Rab5 (B). Comparaison of M1 Manders’ coefficients between SorLA genotypes was performed by the Welch’s t-test, and p-values significances after Bonferroni’s correction are represented above the box plots. For colocalisation analyses, more than ten images were acquired for each co-labelling.

Altogether, these data indicate that the subcellular distribution of the S124R and N371T SorLA variants is similar to the WT form of the protein, in accordance with a wild-type trafficking pattern. In contrast, the three maturation-defective variants (R332W, S577P and R654W) show diverse degrees of subcellular mislocalization, with a smaller proportion of SorLA in early endosomes and a higher proportion of SorLA in the endoplasmic reticulum, consistent with trafficking defects leading to their partial retention in this compartment.

### SorLA variations associated with impaired cellular trafficking result in increased Aβ secretion in hiPSCs

We next tested the impact of the S124R, R332W, N371T, S577P, R654W *SORL1* variants on Aβ secretion. The levels of Aβ40 peptides secreted in the culture media from wild-type, S124R, R332W, N371T, S577P, R654W or KO *SORL1* hiPSCs were measured using the MSD technology (Figure 8, Table S5). As expected, KO *SORL1* hiPSCs resulted in a significant increase in Aβ40 secretion (123.2 ± 14.1 % of control value, p < 0.001). The S124R, N371T and S577P mutants did not show increased Aβ40 secretion compared to wild-type hiPSC (100.4 ± 8.9, 99.1 ± 11.0, and 94.7 ± 12.6, respectively). In contrast, the R332W and R654W mutants, which showed the most obvious effect on SorLA maturation, led to significantly higher Aβ40 secretion (121.4 ± 14.4 p < 0.01 and 127.2 ± 19.6 p < 0.001, respectively).

**Figure 8:**
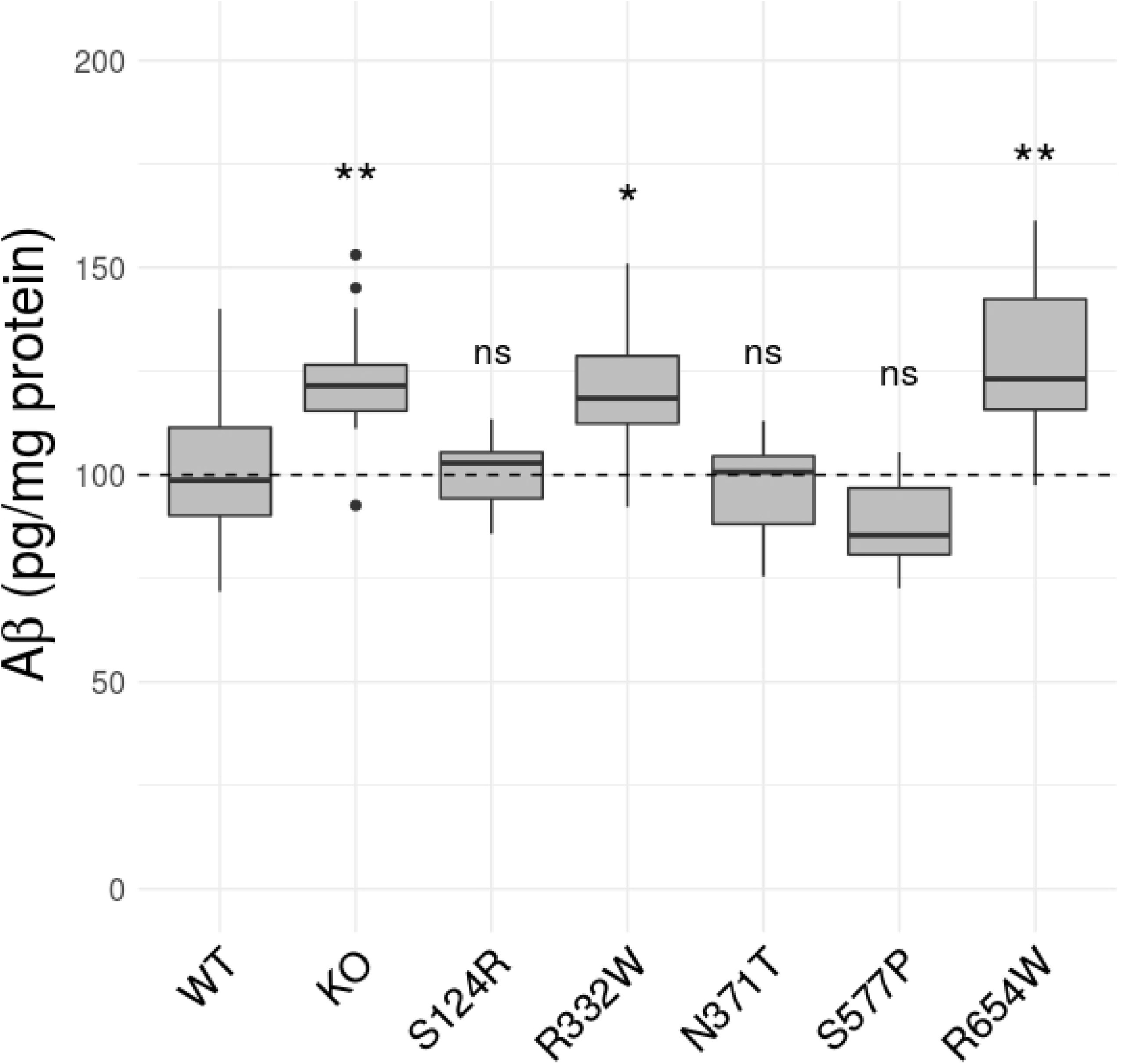
Analysis of Ab secretion by wild-type and S124R, R332W, N371T, S577P, R654W *SORL1* CRISPR/Cas9-edited human iPSC lines. Quantification (pg/mg total protein) of Aβ40 amounts in cell-conditioned medium as determined by MSD assays. Comparisons to WT were performed in one regression model adjusted for both clones and experiments as random effects.

### Impaired cellular trafficking of SorLA protein could be due to subtle variations of the protein 3D structure resulting from changes in the interatomic interactions

The DynaMut webserver (Rodrigues *et al*, 2018) was used to evaluate the impact of missense variations on SorLA structural stability. We focused on variations localized in the Vps10p domain of SorLA whose crystal structure was available in the Protein Data Bank (Kitago et al., 2015, PDB ID 3WSY). Both variants corresponding to maturation-defective proteins (S114R, R332W, G543E, S564G, S577P, R654W and R729W) or displaying a similar maturation profile as the one observed for WT-SorLA (S124R, D140N, Y141C, E270K, N371T, C473S, R490L, G511R, S602L and N674S) exhibited similar profiles (Table S2). Moreover, the variations in the Gibbs Free Energy values was relatively small, suggesting that the mutations had no major impact on SorLA global structure.

To gain further insight into the effect of missense variations on SorLA function, amino acids associated with variants were displayed on the protein structure (Figure 9). The 10 variations presenting a wild-type maturation profile corresponded to exposed amino acids as expected for neutral variations (Shanthirabalan *et al*, 2018). Positions corresponding to impaired maturation variants were distributed over the Vps10p domain. The associated residues were either buried (G543E) or partially buried (S564G, S577P, R654W, Figures 9C and 9D) or partially exposed (S114R, R332W, R729W, Figures 9A and 9B) but did not correspond to core residues. These observations were consistent with the lack of major effects on the protein structure stability predicted by DynaMut and indicated that the altered function was rather linked to modifications of macromolecular interactions such as hydrophobic and electrostatic interactions, van der Waals forces and hydrogen bonds.

**Figure 9:**
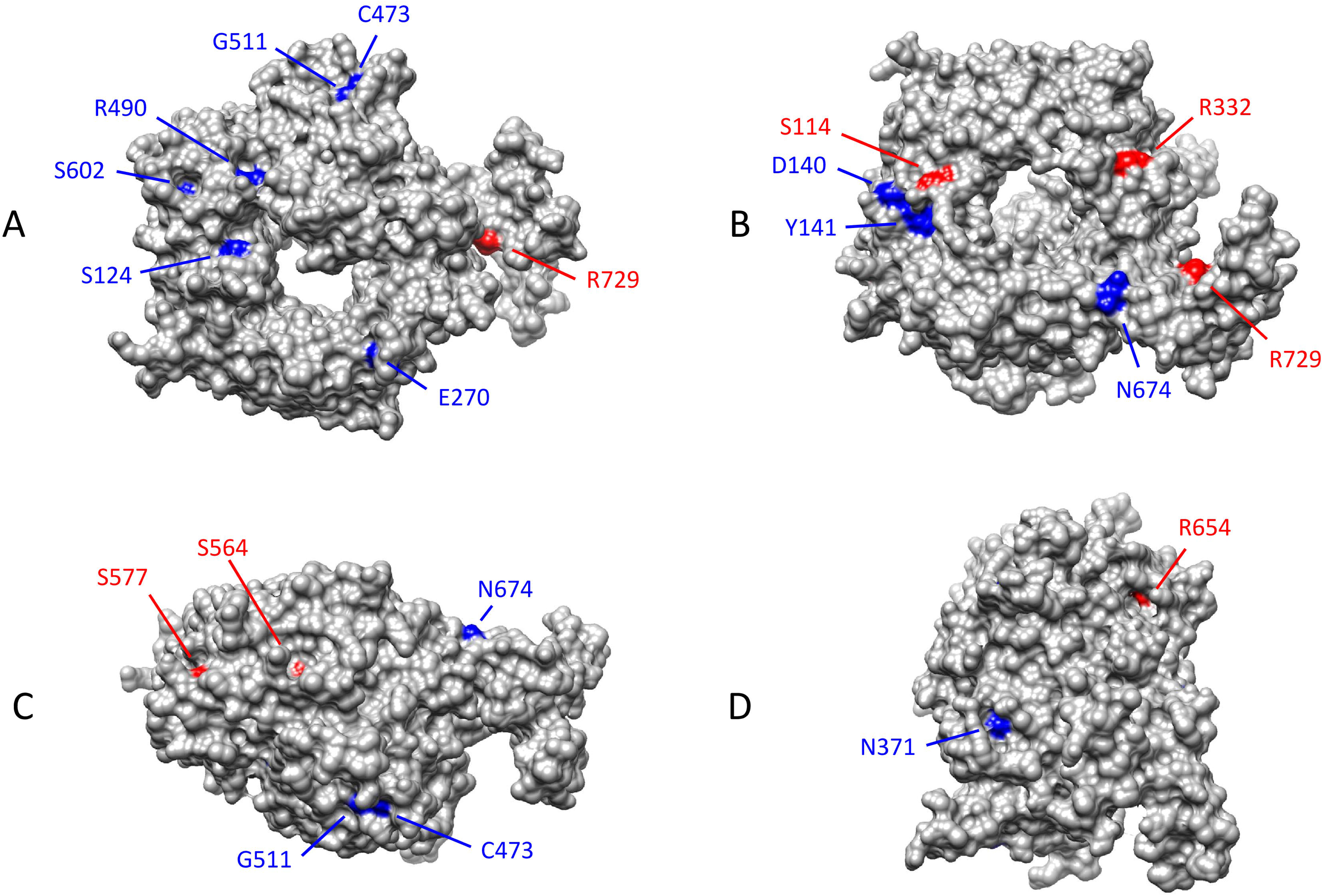
Representation of the amino acids associated with missense variations on the molecular surface of the wild-type human SorLA Vps10p domain (PDB ID 3WSY). Positions associated with impaired maturation are colored in red, those which have no effect are colored in blue. (A) Top view; (B) Bottom View; (C and D) Rotation by 90° around the vertical or horizontal axes of (A) representation. The G543E variation, which is totally buried, is not observable.

The interatomic interactions of the 3 impaired maturation and deficit-trafficking variants (R332W, S577P, and R654W) and of the 2 maturation-competent variants (S124R and N371T) studied in hiPSC were further studied in detail (Figure 10). Consistent with its neutral nature, the mutation of a Serine to an Arginine in the S124R variant, despite the size difference between the two residues, led to very few changes in the intermolecular interactions (Figures 10A and 10A’). Similarly, for the N371T variant, the interaction network, except a hydrogen bond between N371 and E388, was conserved (Figures 10C and 10C’). On the contrary, for the last three variants (R332W, S577P, and R654W), the interaction network as well as the contact nature were modified. In the R332W variant, the substitution of a long polar by a bulky aromatic sidechain gave rise to additional contacts, such as cation-π interactions with R279 and to the loss of the hydrogen bond with D304 (Figures 10B and 10B’). Other existing interactions were altered, going from polar or ionic to hydrophobic. In a similar manner, for S577P, the replacement of a polar by a hydrophobic sidechain, changed the nature of the interaction network, polar contacts being switched for hydrophobic ones (Figures 10D and 10D’). As for R332W, in the R654W variant, the substitution of an Arginine by a Tryptophane led to the appearance of cation-π interactions with G547 and of hydrophobic contacts with E567, as well as to the disappearance of the H-bond with G547 (Figures 10E and 10E’). Altogether, these analyses suggest that altered maturation and trafficking of SorLA protein could be due to subtle variations of the protein structure resulting from changes in the interatomic interactions.

**Figure 10:**
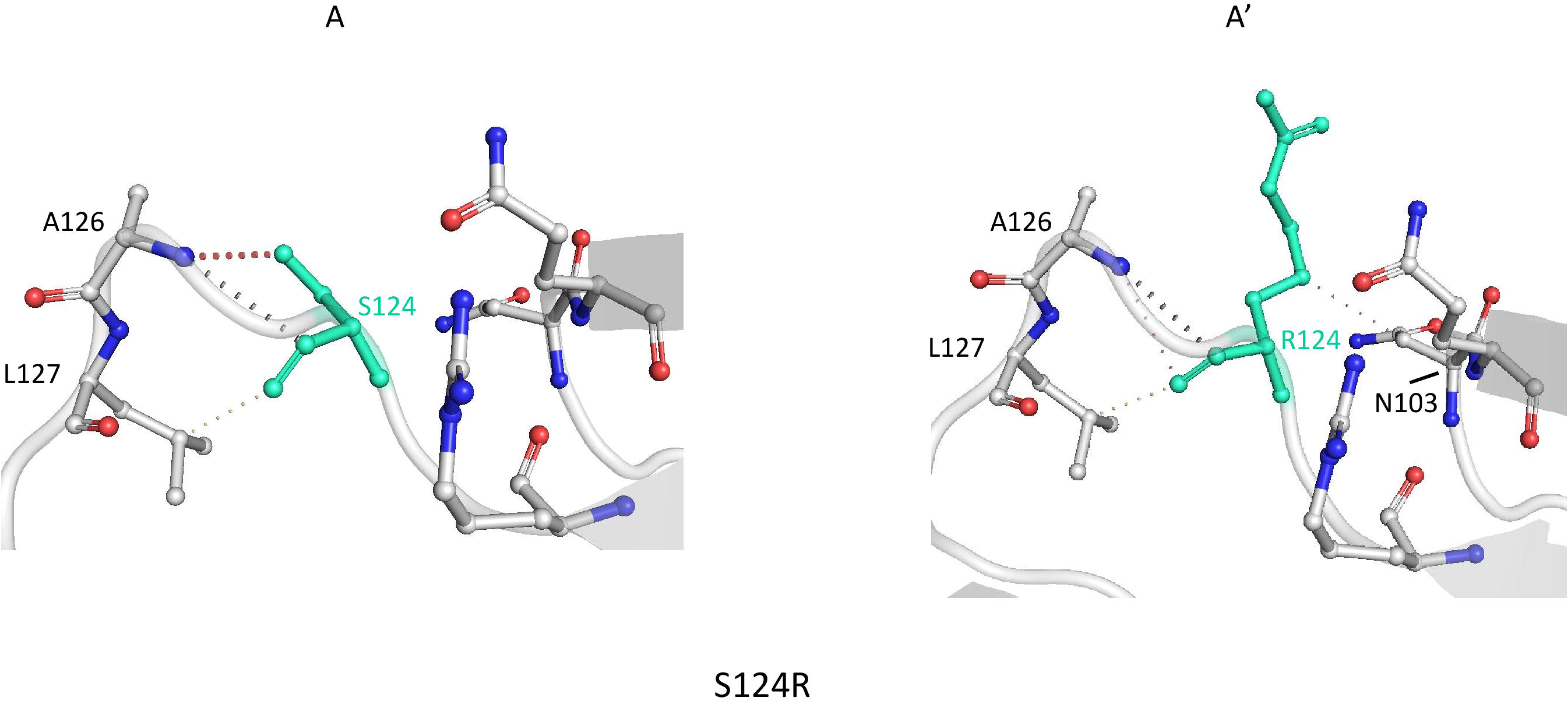
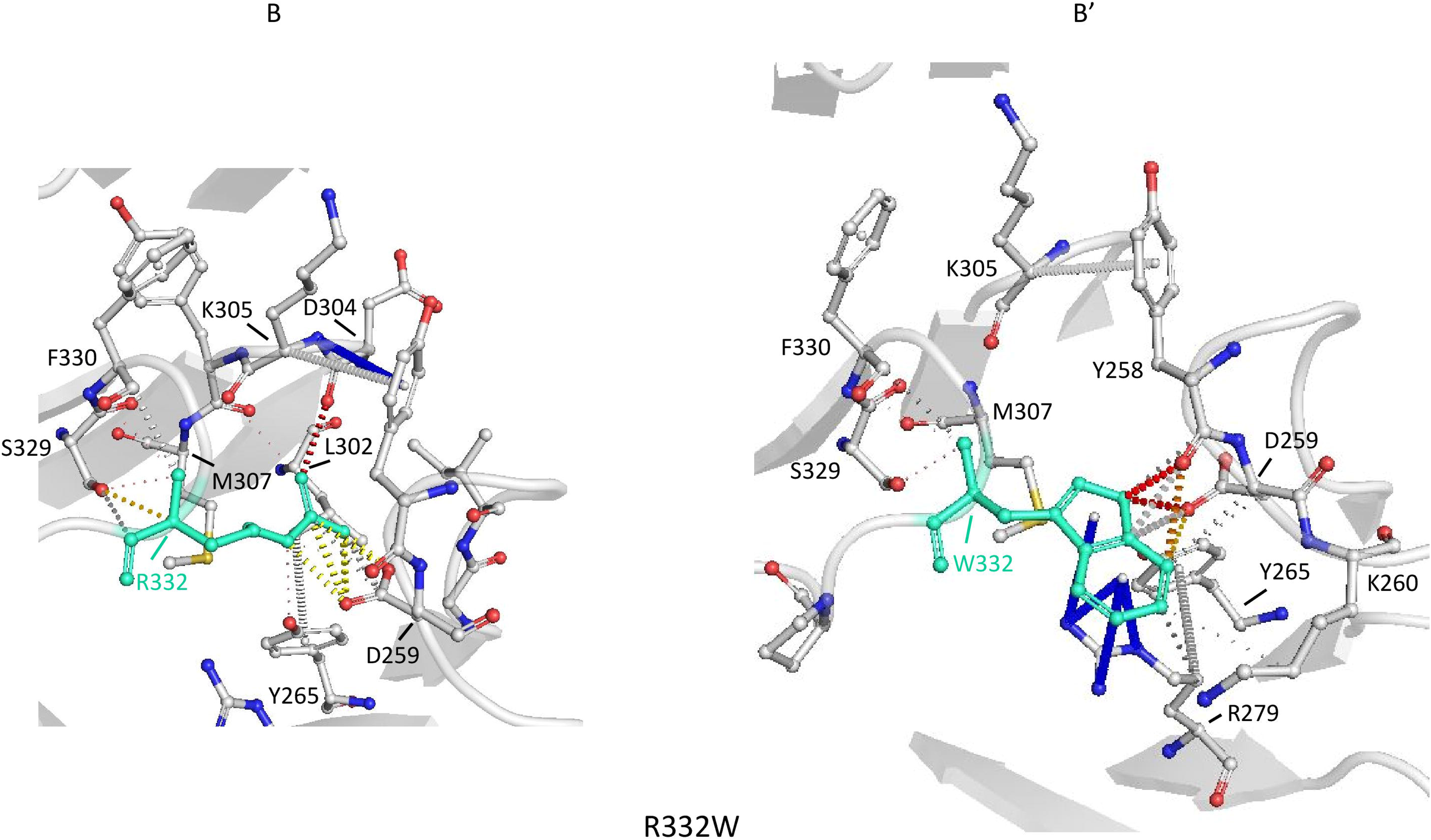
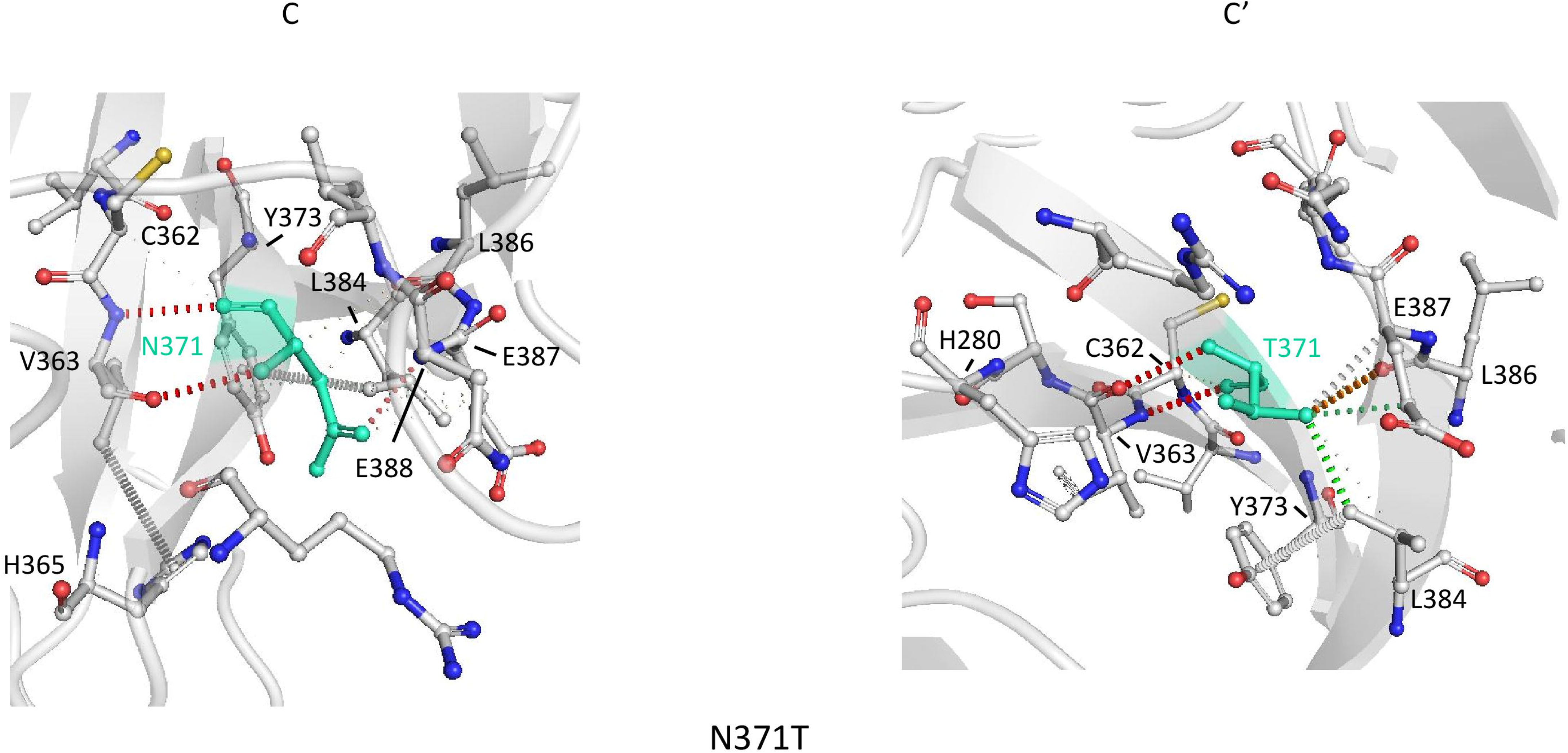
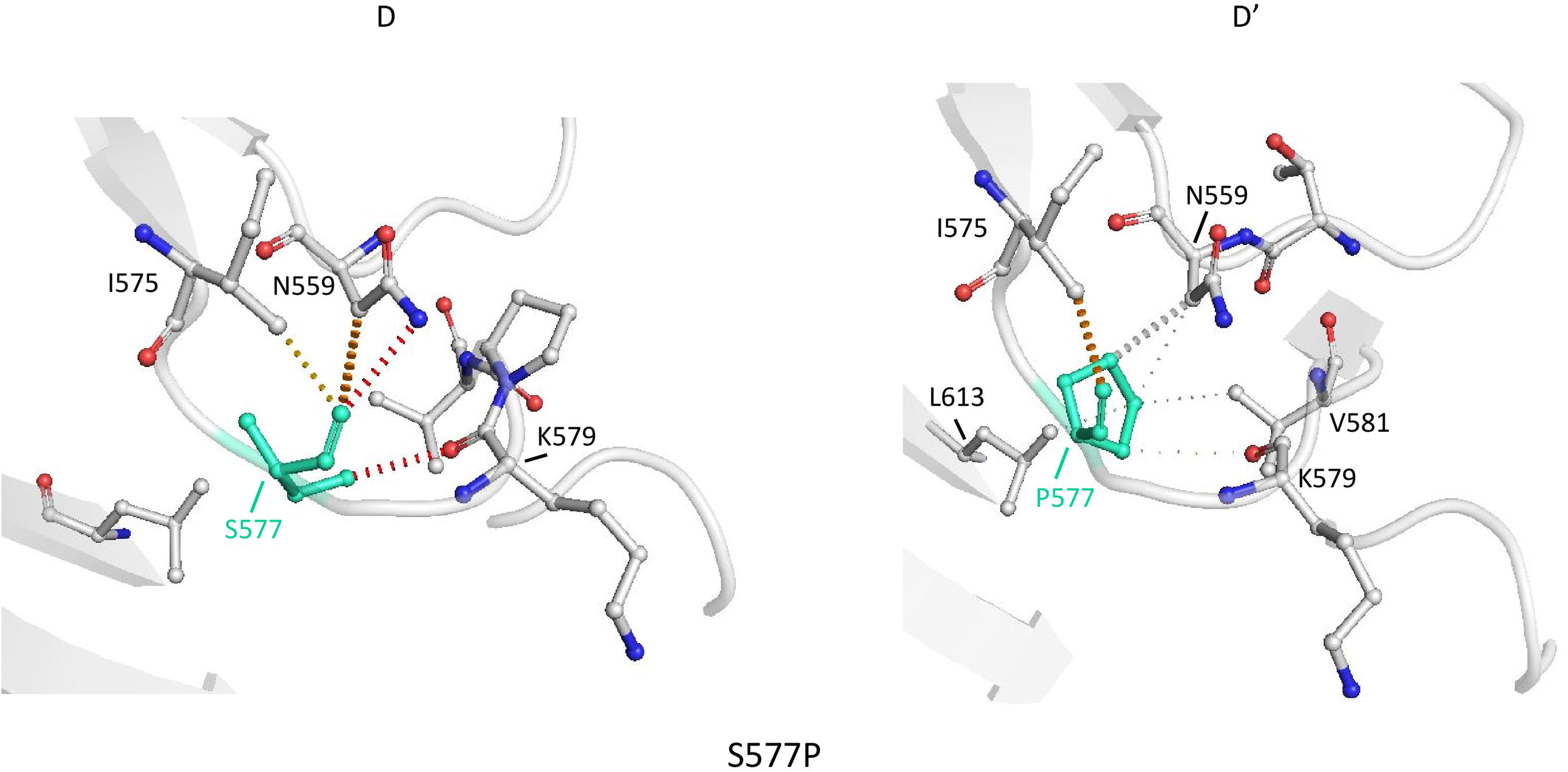
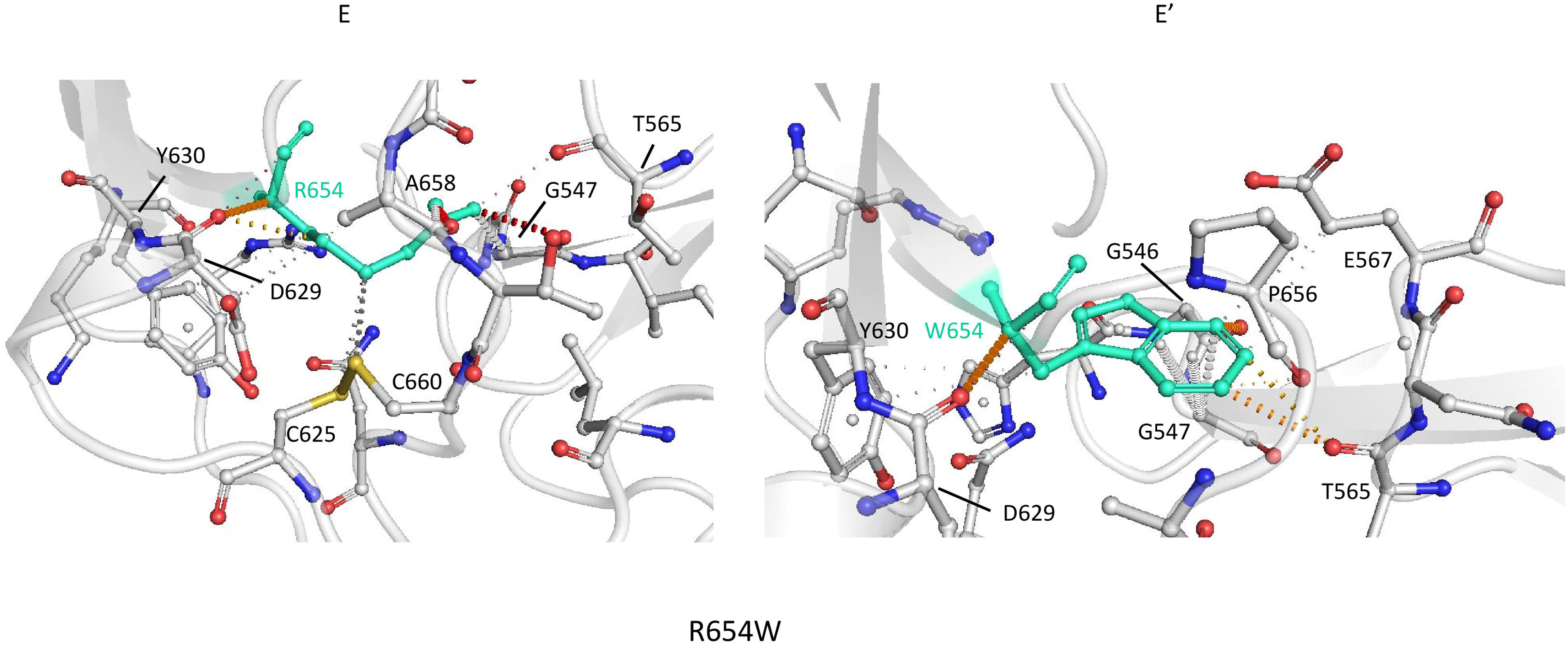
Prediction of interactions between amino acid residues in human SorLA Vps10p domain (PDB ID 3WSY) by DynaMut. The WT and variant residues are colored in green and are represented as sticks alongside the surrounding residues that are involved in any type of interactions. Amino acids involved in interactions are labelled in black and labels are placed next to Cα atoms. The colors of the contacts are based on the following: red, polar (including hydrogen bonds); orange, weak polar; yellow, mixed ionic van der waals; blue, cation π; and gray, hydrophobic (including carbon π). Interactions between the local amino acids of the WT and (A and A′) the S124R variant; (B and B′) the R332W variant; the N371T variant (C and C′); the S577P variant (D and D′); the R654W variant (E and E′).

## DISCUSSION

Previous functional studies proposed two distinct pathological molecular mechanisms for *SORL1* misense variants: (i) reduced binding affinities for APP and thus increased amyloidogenic processing of APP, by favouring its targeting to the late endosome compartments where APP is cleaved into Aβ, or (ii) reduced Aβ binding that may affect the ability of SorLA to direct Aβ peptides to lysosomes for degradation (Caglayan *et al*, 2014; Vardarajan *et al*, 2015; Cuccaro *et al*, 2016). In the present study, we identified a novel pathological molecular mechanism, in which rare missense SorLA variants show protein maturation and trafficking defects.

We analyzed 71 rare missense variants identified from exome sequencing in AD patients by our group. When overexpressed in HEK293 cells, 15 of these 71 variants (S114R, R332W, G543E, S564G, S577P, R654W, R729W, D806N, Y934C, D1535N, D1545E, P1654L, Y1816C, W1862C, P1914S) showed a maturation and trafficking-deficient phenotype. Five of these variations (S124R, R332W, N371T, S577P, and R654W) were further studied in details in CRISPR/Cas9-modified hiPSCs in order to avoid any bias related to the overexpression of the SorLA protein. When expressed at endogenous levels, the maturation defective profile of R332W, S577P, and R654W SorLA variants was confirmed. We further demonstrated that these variants were largely retained in the endoplasmic reticulum, explaining the higher content of high mannose type *N*-glycans observed. This resulted in a reduction in the delivery of SorLA mature protein to the plasma membrane and to the endosomal system. Importantly, this maturation/trafficking-defective process was associated with a clear increase of Aβ secretion for the R332W and R654W variants, demonstrating a loss-of-function effect of these SorLA variants regarding this ultimate readout, and a direct link with AD pathophysiology. Futhermore, structural analysis of the impact of missense variations on SorLA protein revealed that impaired cellular trafficking of SorLA could be due to subtle variations of the protein structure resulting from changes in the interatomic interactions.

Defects in protein trafficking and ER retention have been shown to be a common cellular mechanism involved in the pathogenesis of a wide range of inherited human diseases, such as cystic fibrosis (Welsh & Smith, 1993) or kidney diseases (Schaeffer *et al*, 2014). Mutations act by impacting protein folding and conformational stability, thus interfering with efficient post-translational modification and subsequent protein trafficking. The transport of misfolded glycoproteins in the ER is inhibited by a process called quality control (Adams *et al*, 2019). Pending the severity of the conformational defect, partially folded proteins can be chaperoned to attain their native conformation whereas irreversibly misfolded polypeptides are disposed via the ER-associated degradation pathway (ERAD).

Our strategy, based on an initial screening in HEK293 cells followed by a subsequent validation in CRIPSR/Cas9-modified hiPSCs, revealed a few discrepancies between our cellular models. The effect of the R332W and S577P variations on SorLA maturation profile, which was similar to that of the R654W variant in HEK293 cells, appeared to be weaker in hiPSCs. Indeed, some mature forms of R332W and S577P SorLA proteins were produced and addressed to the plasma membrane in hiPSCs. Overexpression may therefore exacerbate the effect of some variants. HEK293 cells overexpressing SorLA would contain such large amounts of the protein that the machinery responsible for further post-translational processing and the ER quality-control system may well be overloaded, leading to a disproportionate accumulation of immature forms. The HEK293 cells might thus be a sensitive model to reveal variations with low or moderate impact on SorLA maturation.

Cell-surface biotinylation experiments in CRISPR/Cas9-modified hiPSCs showed that, when expressed at endogenous levels in hiPSCs, only the glycosylated mature form of the SorLA protein was detected at the plasma membrane, suggesting a prerequisite of SorLA maturation for its proper trafficking to the plasma membrane. Interestingly, although at very low level, the R654W variant exists at the cell membrane in two forms: an immature and a mature form. ER-retained proteins are typically degraded in a pre-Golgi compartement. However, in some diseases, it has been shown that ER-retained mutant proteins are not effectively degraded and might accumulate, which seems to be also the case for the immature R654W mutant protein. Thus, when present in a great amount in the ER, some immature species and/or misfolded protein might bypass the ER quality control mechanism and be transported to the plasma membrane. These data are fully consistent with our results obtained in HEK293 cells in which we detect a disproportionate accumulation of immature forms, and the presence of both immature and mature forms at the plasma membrane. The presence of both species at the plasma membrane in HEK293 cells, which is consistent with the data previously reported by other groups (Christensen *et al*, 2020; Jacobsen *et al*, 2001), could therefore be related to the overexpression of SorLA and the saturation of the ER quality control system.

The 15 defective-trafficking SorLA variants identified in this study (S114R, R332W, G543E, S564G, S577P, R654W, R729W, D806N, Y934C, D1535N, D1545E, P1654L, Y1816C, W1862C, P1914S) are distributed along the entire extracellular part of the protein, with the exception of the EGF-like and the LDLR class A repeats domains, suggesting that these two domains are less sensitive to amino acid substitutions regarding this particular phenotype. We wondered how these 15 amino acid substitutions could impact SorLA protein global structure, stability and/or flexibility. None involves an Asparagine (N) potentially involved in N-glycosylation anchoring. Nor do they create ER-resident motifs, such as KDEL and dilysine. The Y1816C and W1862C variations that lead to the introduction of additional Cysteine residues within the Fibronectin type III-cluster may be involved in the formation of incorrect disulfide bonds within the domain or even other intramolecular disulfide bonds. The analysis of the impact of missense variations on SorLA structural stability, focusing in the Vps10p and a part of the SortilinC domains of SorLA whose crystal structure was available (Kitago *et al*, 2015), suggested that the mutations had no major impact on SorLA global structure. However, the detailed study of interatomic interactions of the 3 impaired maturation variants (R332W, S577P, and R654W) showed that the interaction network as well as the contact nature were modified. Alteration of the interatomic interactions could result in subtle variations of the protein 3D structure and folding stability.

The study in hiPSCs allowed us to highlight a gradual effect of the variants, with S577P<R332W<R654W. This gradual effect could be observed on all the phenotypes assessed: maturation defects of the protein, retention in the ER, release to the plasma membrane, endosomal localization and Aβ production. This could be due to different severities of the folding defect present in the mutated protein and/or the ability of the ER quality control system to correct these. We can assume that the impact of the R332W and S577P variations on protein conformation in this cellular model are more moderate, and that the folding process is able to occur, at least partially.

In contrast to the R654W variant that was barely detected as mature species, significant amounts of mature forms of R332W and S577P SorLA mutant proteins were produced in hiPSCs, raising the question of their functionality regarding APP metabolism. The expression of the R332W variant resulted in a significant increase of Aβ secretion. This effect on extracellular Aβ levels may result from an insufficient amount of mature SorLA protein in the appropriate cellular compartments to ensure its function with respect to APP metabolism, and/or a direct impact of this variation on SorLA function on APP metabolism, both not being exclusive. Regarding the S577P variant, we observed no effect on extracellular Aβ levels. This indicates that, in our cellular model, the amount of mature protein produced is sufficient and that this mutant protein is functional regarding APP metabolism. The absence of effect of the S577P variant on Aβ secretion in hiPSCs does not preclude its role in the AD neurodegenerative process. Indeed, age-associated deterioration of cellular machinery leads to an increase in the occurrence of protein misfolding, accumulation and aggregation, due in part to the gradual decay of ER chaperoning systems. It is therefore possible that this S577P variant, identified in AD patients, has a long-term effect that would not be featured by our short-term culture experiments. As we cannot exclude specific effects of SorLA variants during aging conditions, neither can we exclude a cell-specific effect. It would be interesting to model this variant, and also the S124R and N371T variation for which we detected no effet effect on Aβ secretion, in a neuronal environment with long-lasting experiments to determine whether they are involved in AD or if they should be considered as rare sequence variants with no functional significance and no relevance with AD.

Interestingly, genomic data appear to be in line with our *in vitro* experiments, since we observed an apparent inverse correlation between the frequency of the variants and the measured effect in our experiments. Indeed, the frequency in large control databases such as gnomAD v2.1.1, non-neuro subset, ranged from a full absence (R654W), to the presence in 4 individuals (R332W), 9 indididuals (S577P) and 304 individuals (N371T), out of 114,679 controls. This is consistent with the inverse correlation between odds ratios and variant frequency found in case-control studies (Campion *et al*, 2019; Holstege *et al*, 2017). Conversely, the S124R variant, which is recurrent among AD cases (2 French cases, through two distinct genomic changes), is absent in controls and in the gnomAD database. Thus, the existence of two distinct genomic changes leading to a same missense change in two unrelated patients suggests that this amino acid position should be further studied in other models, as for the S577P, and despite the absence of modification of Aβ secretion in hiPSCs after 24h secretion.

Of note, case-control studies assessing the putative effect of *SORL1* missense variants mainly relied on the use of *in silico* predictions (e.g., the Mis1-2-3 classification). Interestingly, all 15 variants showing an impact on SorLA maturation in HEK293 cells belonged to the Mis3 category. However, such *in silico* predictions are not a sufficient argument for using *SORL1* variants in a clinical context, as not all Mis3 variants might eventually behave as loss-of-function variants, first, and because there may be some diversity in the types and strengths of effects among missense variants, second.

In conclusion, by assessing missense variants located throughout the coding sequence of the *SORL1* gene among rare variants found in Alzheimer disease patients, we identified a novel mechanism leading to a loss of SorLA function. A subset of the variants induced a likely misfolded protein, thus altering the protein maturation and trafficking, and eventually leading to a loss of the protective function of SorLA towards Aβ secretion. *SORL1* maturation defective variants are a novel mechanism directly link to AD pathophysiological mechanisms.

## MATERIALS AND METHODS

### cDNA constructs

The SorLA^759^ and SorLA^2131^constructs were designed following the constructs described in (Jacobsen *et al*, 2001). The wild-type human SorLA^759^ (Met^1^-Glu^759^) cDNA construct was generated by PCR-amplification on the pCMV6-XL5-human full-length SORL1 cDNA plasmid (pCMV6-XL5-WT-SorLA^FL^ OriGene Technologies, Inc, Rockville, MD, USA) using the 5’ CCGGAATTCCGGCAAAATGGCGACACGGAGCAGCAGG 3’ and 5’ TGCTCTAGAGCACTACTCGTTCTCTTCTGCCAGGGG 3’ oligonucleotides. Residues number refer to the primary structure of the wild-type protein. The SorLA^759^ PCR products was next subcloned as an EcoRI/Xba fragment into the pCMV6-XL5 expression vector. The wild-type human SorLA^2131^ (Met^1^-Ala^2131^/Val^2209^-Ala^2214^) cDNA construct was generated using the GeneArt™ Site-Directed Mutagenesis System (Invitrogen^TM^, Carlsbad, CA, USA) and a PCR overlap extension procedure on the pCMV6-XL5-human full-length SORL1 cDNA plasmid. The sequences of mutagenized oligonucleotides were: 5’ GCA TCT GCA ACG CAG GCT GCC GTC CCC ATG GTG ATA GCC TGA AAG AGC 3’ and 5’ GCT CTT TCA GGC TAT CAC CAT GGG GAC GGC AGC CTG CGT TGC AGA TGC 3’. Integrity of the SorLA^759^ and SorLA^2131^ cDNA constructs were verified by sequencing. Site-directed mutagenesis was then carried out over the pCMV6-XL5-WT-SorLA^FL,^ pCMV6-XL5-WT-SorLA^759^ or pCMV6-XL5-WT-SorLA^2131^ plasmids using the QuikChange II XL Site-Directed Mutagenesis Kit from Agilent (Santa Clara, CA, USA) according to manufactory’s instructions. All oligonucleotides used were listed in Table S3. Integration of the mutation were verified by sequencing. The plasmidic constructs with S114R, R332W, G543E, S564G, S577P, R654W, R729W, D806N, Y934C, D1535N, D1545E, P1654L, Y1816C, W1862C, P1914S SorLA variations were sequenced in their entirety.

### HEK293 cells culture and transfection

HEK293 cells were grown and maintained in DMEM/F12 medium (Gibco/Thermo Fisher Scientific, Waltham, MA, USA) supplemented with 10% FCS (Eurobio, Les Ulis, France). Cells were plated in 12-well or 6-well plates 48 h prior to transfection, grown to approximately 90% confluence and transfected with the appropriate amounts of the indicated constructs by using the lipofectamine 3000 reagent (Invitrogen^TM^) according to the manufacturer’s protocol. Twenty-four or 48 hours after transfection, the cells were harvested and processed for Western blotting or biotinylation analyses.

### hiPSC culture

The hiPS cell line was previously generated by our group. This cell line derives from an unaffected male donor and has an APOE 3/3 genotype. The line was analyzed for pluripotency markers expression by quantitative PCR and for chromosomal abnormalities by karyotyping and was able to differentiate into the three germ layers. Whole exome sequencing of the parental hiPSC line confirmed the absence of rare coding variants in the *SORL1* gene, as well as in *APP, PSEN1, PSEN2, TREM2, ABCA7* and other genes causing Mendelian forms of dementia. hiPSCs were cultured on feeder-free conditions in mTeSR Plus medium (STEMCELL Technologies, Vancouver, Canada) on Matrigel-coated culture dishes (Corning, Corning, NY, USA) diluted in DMEM-F12 according to manufacturer’s instructions) in a 37°C/5% CO_2_ incubator. Cells were split when they reached 80% confluency using StemPro Accutase (Thermo Fisher Scientific) and plated in 10µM ROCK inhibitor (StemGent, Cambridge, MA, USA) supplemented medium. Differentiating colonies were removed from the plate before splitting. Cell lines were confirmed to be free of mycoplasma.

### CRISPR/Cas9 genome editing

Genome editing was performed following a published protocol (Ran *et al*, 2013). The guide RNAs (gRNAs) were designed using the CRISPOR.org web tool (Concordet & Haeussler, 2018) (http://crispor.tefor.net/). Two gRNAs were designed for each variant. We chose guide sequences located as close to the mutation site, with the highest specificity score (Table S4). All gRNAs were cloned into the pX458 vector (Addgene, pSpCas9(BB)-2A-GFP, plasmid #48138) that coexpresses the Cas9 nuclease and the GFP. Each plasmid and the corresponding single-stranded oligodeoxynucleotides (ssODN) were nucleofected in the hiPSC line, using an AMAXA nucleofector II device. Two days later, cells were stained using DAPI (NucBlue^TM^ Fixed Cell ReadyProbes reagent, Molecular Probes, Thermo Fisher Scientific) as viability dye. GFP-positive cells among living cells were sorted and plated as 1 cell/well into 96-well plates using FACS Aria III (BD Bioscience, Franklin Lakes, NJ, USA). When clones reached 80% confluency, genomic DNA was isolated and analysed by Sanger sequencing (Table S4). For each variation, 6 clones were selected, including three clones carrying the variation at the homozygous state and three isogenic wild-type clones. For *SORL1 KO-/-* selection, we selected clones carrying a homozygous frameshift insertion. The absence of the protein was then verified using Western blotting (Fig S6).

### Biotinylation

Subconfluent cultures were rinsed twice with PBS+/+, exposed to sulfo-NHS-SS-biotin (Thermo Fisher Scientific) for 30 min at 4°C, rinsed three times with glycine quenching buffer (100 mM glycine in PBS+/+) and solubilized in RIPA buffer (50 mM Tris/HCl pH8, 150 mM NaCl, 0.5% (w/v) deoxycholic acid, 1% NP40, 10% glycerol, 2mM DTT), supplemented with a cocktail of protease inhibitors (Sigma-Aldrich) and phosphatase inhibitors (Halt phosphatase, Thermo Fisher Scientific) for 10 min on ice. The resulting lysate was centrifuged at 12,000 x g for 10 min at 4 °C, and total cellular protein content was determined using the DC Protein Assay Kit (Bio-Rad Laboratories, Hercules, CA, USA). Then, 200 µg of proteins were incubated with streptavidin beads (Thermo Fisher Scientific) for 1h at 4 °C (4 μg of protein/μl of beads pre-equilibrated with RIPA buffer). The supernatant was removed using a magnetic stand, then beads were washed five times with RIPA buffer, and resuspended with 1× Laemmli sample buffer (Bio-Rad Laboratories).

### Protein extraction

Cells were homogenized in RIPA buffer (Tris-HCl pH8 0.05M, NaCl 0.15M, NP-40 1%, Sodium deoxycholate 0.5%, Glycerol 10%, DTT 2mM), supplemented with a cocktail of protease inhibitors (Sigma-Aldrich, Saint-Louis, MI, USA) and phosphatase inhibitors (Halt phosphatase, Thermo Fisher Scientific). After 10 min on ice, lysates were centrifuged (12,000 × g, 10 min, 4°C) and the supernatant containing soluble proteins was collected. Protein concentrations were measured using the DC Protein Assay Kit (Bio-Rad Laboratories).

### Glycosidase treatments

PNGase F, O-glycosidase, Neuraminidase and Endo H were obtained from New England Biolabs (Ipswich, MA, USA). Digestions were performed according to the manufacturer’s instructions. All digestions were carried out for 24 h at 37°C for PNGase F/O-glycosidase/ neuraminidase treatments and for 3h for Endo H treatments. After this, the proteins were analyzed by SDS-PAGE as below.

### Western blotting

Proteins were resolved by Tris-acetate NOVEX NuPAGE 3-8% (Invitrogen) (SorLA^FL^ and SorLA^2131^) or TGX Stain-Free gels (Bio-Rad Laboratories) (SorLA^759^ and samples from biotinylation experiments) then transferred onto nitrocellulose membrane using the Trans-Blot Turbo system (Bio-Rad Laboratories). Membranes were then blocked in 5% non-fat milk and immunoblotted with appropriate primary antibodies: mouse monoclonal anti-SorLA (1:5,000; mAb# 611860, BD Bioscience) for the detection of full length SorLA and SorLA^2131^ proteins, rabbit monoclonal anti-SorLA antibody specific to the N-terminus of human SorLA protein (1:1,000; D8D4G #79322; Cell Signaling, Danvers, MA, USA) for the detection of SorLA^759^ proteins, rabbit polyclonal anti-FUS (1:5,000; A300-302A; Bethyl Laboratories, Montgomery, TX, USA). Membranes were then incubated with secondary peroxidase-labelled anti-mouse or anti-rabbit antibodies (1:10,000) from Jackson Immunoresearch Laboratories (WestGrove, PA, USA), and signals were detected with chemiluminescence reagents (ECL Clarity, Bio-Rad Laboratories). Signals were acquired with a GBOX (Syngene, Cambridge, UK), monitored by the Gene Snap software (Syngene). When appropriate, the signal intensity in each lane was quantified using the Genetools software (Syngene) and normalized with the Stain-Free signal quantified in the corresponding lane (ImageLab™ software, Bio-Rad Laboratories).

### Aβ secretion assay

HiPSCs were plated on matrigel pre-coated 12-well plates in triplicate. Two days later, medium was replaced by 500µL of mTeSR Plus supplemented with phosphoramidon (10µM, Sigma-Aldrich). Twenty-four hours later, conditioned medium was collected in each well and centrifuged at 1,000 x g for 5 minutes. The supernatant was supplemented with a cocktail of protease inhibitors (Sigma-Aldrich) and frozen at −80°C until use. The quantification of Aβ peptides was performed using the V-PLEX Aβ Peptide Panel 1 (6E10) kit (Meso Scale Diagnostics, Rockville, MD, USA) according to manufacturer’s instructions. Conditioned media were diluted 1:2 in Diluent 35 and analyzed in duplicate. The plate was analyzed using a MESO QuickPlex SQ 120 instrument (Meso Scale Diagnostics). Aβ quantities were normalized to the total protein amount measured in each sample.

### Immunofluorescence

HiPSCs were plated on matrigel-coated 20 mm round coverslips. Twenty-four hours later, cells were rinsed twice with PBS before fixation with 1 ml of 4% paraformaldehyde (PFA) for 1 hr at room temperature followed by three washes in PBS of 10 min each. Antigen retrieval was performed according to Hayashi et al. (Hayashi *et al*, 2011). Briefly, coverslips were immersed in a 6M urea - 0.1M Tris (pH9.5) bath at 80°C for 10 min, followed by three washes in PBS of 15 min each. Cells were permeabilized with PBS/0.1% Triton X-100 for 5 min, and blocked for 30 min in antibody buffer [PBS containing 2% bovine serum albumin (BSA) and 3% normal goat serum (NGS)]. Cells were then incubated with primary antibodies in antibody buffer for 1h at room temperature. Following three washes with PBS (10 min each), cells were incubated with secondary antibodies in antibody buffer at room temperature for 1 hrin dark. Following two washes with PBS (10 min each), cells were counterstained with DAPI (NucBlue^TM^ Fixed Cell ReadyProbes reagent, Molecular Probes, Thermo Fisher Scientific) and coverslips were finally mounted in ProLongTM Diamond antifade solution (Thermo Fisher Scientific). The following antibodies were used: anti-SorLA 48/LR11 (1:200; 612633; BD Biosciences), anti-Rab5 (1:250; ab218624; Abcam, Cambridge, UK), anti-Cyclophilin B (1:500; ab16045; Abcam), goat anti-mouse IgG Alexa Fluor (1:600; Molecular probes, Thermo Fisher Scientific), goat anti-rabbit IgG Alexa Fluor (1:600; Molecular Probes, Thermo Fisher Scientific). Images were acquired using a Leica THUNDER Imaging System (Leica Microsystems, Wetzlar, Germany), combining Computational Clearing with Large Volume adaptative deconvolution (LVCC) for post-acquisition processing. Brightness and contrast of images were adjusted using Fiji software (Schindelin *et al*, 2012).

### Co-localization analysis

Co-localization analysis of SorLA with endoplasmic reticulum and early endosomes markers was performed using JACoP plugin (Bolte & Cordelières, 2006) on Fiji environment (RRID:SCR_002285) and presented as M1 Manders’ correlation coefficients. The Manders’ M1 coefficients indicate the proportion of SorLA (green channel) coincident with Rab5 or Cyclophilin B (red channel) over its total intensity.

### Structural analysis

The structure of the VPS10P domain of SorLA in complex with its own propeptide fragment (PDB ID 3WSY) was used as a model for structural analysis (Kitago *et al*, 2015). The changes of stability and flexibility of the VPS10P domain upon mutation were estimated with the DynaMut webserver (Rodrigues *et al*, 2018) (http://biosig.unimelb.edu.au/dynamut/). The structures were analyzed using PyMOL (The PyMOL Molecular Graphics System, Schrodinger, LLC) or UCSF Chimera (Pettersen *et al*, 2004) softwares.

### Statistical analysis

For each experiment, we compared the variants to the WT forms of SorLA by performing a regression model. Variants were all considered into one independent factor where the WT was the reference level. In Western blot experiments, the outcome was the normalized signal intensity, in log scale to overcome heteroscedasticity whereas in Aβ assay, the outcome was the normalized quantification of Aβ peptide. Models were all adjusted for experiment settings. For each variant, the p-value refers to the nullity test of its associated coefficient in the model. We adjusted p-values for multiple testing using the Bonferroni’s correction. Statistical models were performed using the R software. All tests were two-tailed at the significance level of 5% (ns: not significant, * p<0.05, ** p<0.01, ***p<0.001).

## DECLARATIONS

### Ethics approval and consent to participate

Not applicable

### Consent for publication

Not applicable

### Availability of data and material

All data generated or analysed during this study are included in this published article [and its supplementary information files]

## Acknowledgements

We thank Pr Muriel Bardor, Laboratoire Glyco-MEV EA 4358, University of Rouen Normandie, Pr Maité Montero, DC2N, University of Rouen Normandie, and Damien Shapman, PRIMACEN University of Rouen Normandie, for helpful discussion. We thank Muriel Bardor for critical reading of the manuscript. Light microscopy was performed at the PRIMACEN imaging platform (Rouen University, France). We thank Cyril Pottier and Kilan Le Guennec for their help on plasmid synthesis. Molecular graphics and analyses were performed with UCSF Chimera, developed by the Resource for Biocomputing, Visualization, and Informatics at the University of California, San Francisco, with support from NIH P41-GM103311.

## Authors’ contributions

Conceptualization; *A.R-L., L.M., S.F., M.L., G.N.;* Formal analysis: *M.L, L.M, A.R-L, C.S., S.F., I.S-M, L.G.;* Funding acquisition: *A.R-L., M.L., G.N., D.C., T.F., I.S-M, L.G.;* Investigation: *A.R-L., L.M., S.F., C.S., S.P., G.R., S.R., L.G., I.S-M., M.L.;* Methodology: *A.R-L., L.M., S.F., C.S., G.R., L.G., I.S-M., M.L.;* Supervision: *M.L., G.N.;* Validation: *C.S., A.R-L., L.M., S.F., S.R., M.L.;* Visualization: *A.R-L., L.M., S.F., C.S., S.R., L.G., I.S-M., M.L.;* Writing – original draft: *A.R-L., L.M., S.F., C.S., L.G., I.S-M., M.L., G.N., D.C*.

---

**Figure S1: Western blot analyses of SorLA^759^ and SorLA^2131^ proteins expressed in HEK293 cells.**

The SorLA^759^ (A) and SorLA^2131^ (B) proteins secreted into the cellular medium (Secreted) and the corresponding cell lysate (Intracellular) were analyzed by immunoblotting using an antibody specific to the N-terminus of human SorLA protein. For each variation, the patterns were confirmed in at least 4 independent replicates. Representative blots are presented. Note that the R654W and R729W variants showed an increase of their steady-state level in the cellular lysates compared to the wild-type SorLA^759^ construct. In most cases, proteins with trafficking defects are degraded by the ERAD/EGAD quality control system, suggesting that these mutant proteins were less effectively degraded by the quality control systems. The molecular masses of marker proteins in kDa are shown on the left.

**Figure S2: Sanger sequencing analysis of CRISPR/Cas9-edited hiPSC**

For each mutation, the electropherogram showing the insertion of the missense point mutation (on the bottom) is compared to the wild-type sequence (on the top). The nucleotide change on cDNA and the corresponding amino acid change on the protein are indicated.

**Figure S3: Maturation defective SorLA variants display reduced cell-surface levels of SorLA protein in CRISPR/Cas9-edited hiPSC**

Surface biotinylation experiments to examine cell-surface levels of SorLA proteins in wild-type or S124R, R332W, N371T, S577P, R654W *SORL1* CRISPR/Cas9-edited hiPSC. Total lysates (Total) and biotinylated fractions (Surface) were analyzed by immunoblotting using an anti-SorLA antibody. FUS, an intracellular protein, was used as an internal control. The absence of FUS in the biotinylated fraction demonstrates the integrity of the plasma membrane during the biotinylation experiment. Representative blots are presented. Immature core-glycosylated and mature complex-glycosylated SorLA are indicated with solid and empty arrowheads, respectively. The molecular masses of marker proteins in kDa are shown on the left.

**Figure S4: Sequencing and protein analysis of *SORL1* KO hiPSCs**

*SORL1* knock-out clones were selected during the screening of the clones carrying the R332W variant (see guide RNAs on Table S3). We chose clones carrying a homozygous indel, each consisting on an insertion of 1 nucleotide. (A) Sanger sequencing of the clones. The nucleotide insertion on cDNA, the corresponding amino acid change on the protein and the position of the premature stop codon are indicated. (B) Western blot showing the absence of SorLA staining in both *SORL1* KO clones, compared to wild-type (WT) clones. For each clone, two independent replicates were analyzed.

**Figure S5: Retention of SorLA maturation defective variants in the endoplasmic reticulum**

Wild-type and S124R, R332W, N371T, S577P, R654W *SORL1* CRISPR/Cas9-edited hiPSC lines were double-labelled against SorLA (in green) and Cyclophilin B (in red). In blue, DAPI counter-staining.

**Figure S6: Decreased level of SorLA maturation defective variants in early endosomes**

Wild-type and S124R, R332W, N371T, S577P, R654W *SORL1* CRISPR/Cas9-edited hiPSC lines were double-labelled against SorLA (in green) and Rab5 (in red). In blue, DAPI counter-staining.

**Figure S7: Colocalization analysis of SorLA maturation defective variants**

(A, B) Scatter plots representing for each SorLA genotype the mean (and its 95% confidence interval, represented by a halo) of linear regressions derived from corresponding two-dimensionnal cytofluorograms (not displayed to gain in visibility). Cytofluorograms depicted the distribution of pixels in two-color images according to their fluorescence intensity in the green channel (SorLA, x-axis) and in the red channel (cyclophilin *B* (A) or Rab5 (B); y-axis). For colocalisation analyses, more than ten images were acquired per co-labelling, leading to the genesis of more than ten cytofluorograms. The color code assigned to each genotype is the same as in Fig. 7. A line coinciding the first bisector indicates a perfect co-localisation. The numbers between brackets refer to the means of the M1 Manders’ coefficients calculated from the corresponding linear regressions. A shift of the linear regression from the first bisector is indicative of a decreased overlap and results in a smaller M1 coefficient. More the regression line deviates from the first bisector, more the overlap will be the M1 coefficient small.

**Table S1: Summary of analyses performed in HEK293 cells expressing wild-type (WT) or missense variants of SorLA, or in CRISPR/Cas9-edited hiPSC**

Nucleotide positions are relative to canonical transcript NM_003105.5. The nucleotide change on cDNA (Base Change), the corresponding amino acid change on the protein (AA Change), and the location along the protein (Functional domain) are indicated. The Mis3, Mis2, and Mis0-1 missense variants examined in this study are shown in red, orange and black, respectively. A (Y) indicates an impact of the variants on SorLA protein maturation, trafficking or function and a (N) indicates no change from the wild-type form of the protein.

**Table S2: Predicted results obtained with the Dynamut Server upon mutation.**

Mutations corresponding to maturation-defective proteins are colored in red. No correlation between impaired maturation and missense variations was observed. Indeed S114R, R332W and R654W were predicted to result in more stable structures (positive ΔΔG) while G543E, S654G, S577P and R729W were associated with a decrease of stability (negative ΔΔG).

**Table S3: Primer pairs used to introduce SorLA genetic variations into the wild-type sequence by site-directed mutagenesis**

**Table S4: Sequences of guide RNAs, ssODNs and primer pairs used for CRISPR/Cas9 gene editing and clone screening.**

For each variation, the nucleotide change (as a function of the genomic position and *SORL1* cDNA) and the resulting protein change are indicated, together with the sequences of the guide RNAs, the corresponding ssODNs and the Sanger sequencing primer pairs. On ssODN sequences, lowercases correspond to intronic sequences. The position of the variant is indicated in red and the nucleotide triplet is underlined. The synonymous variations introduced to preclude re-cutting by the Cas9 are indicated in green and the nucleotide triplet is underlined.

**Table S5: p-Values of statistical analyses**

