## Supplementary figures and images for "Impaired SorLA maturation and trafficking as a new mechanism for *SORL1* missense variants in Alzheimer disease"

### Supplemental Figures

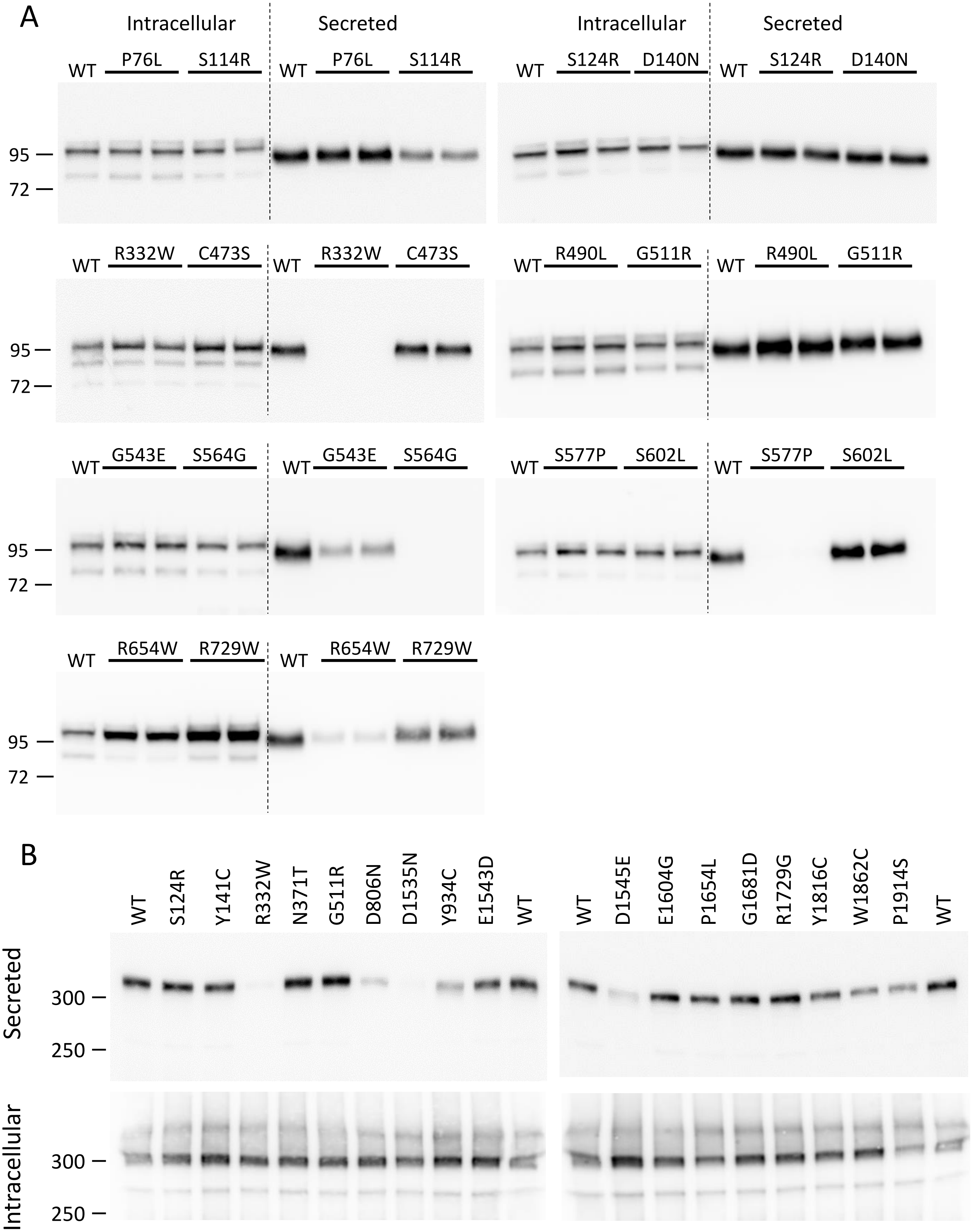

c.372C>G
p.(Ser124Arg)

c.994C>T
p.(Arg332Trp)

c.1112A>C
p.(Asn371Thr)

c.1729T>C
p.(Ser577Pro)

c.1960C>T
p.(Arg654Trp)

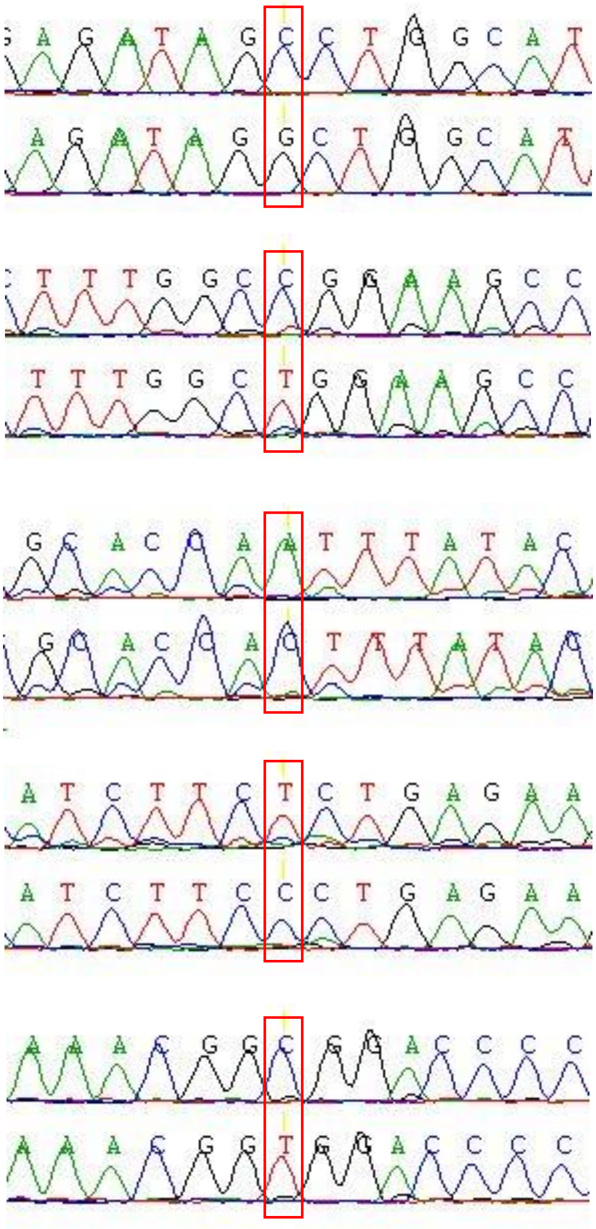

Fig.S2

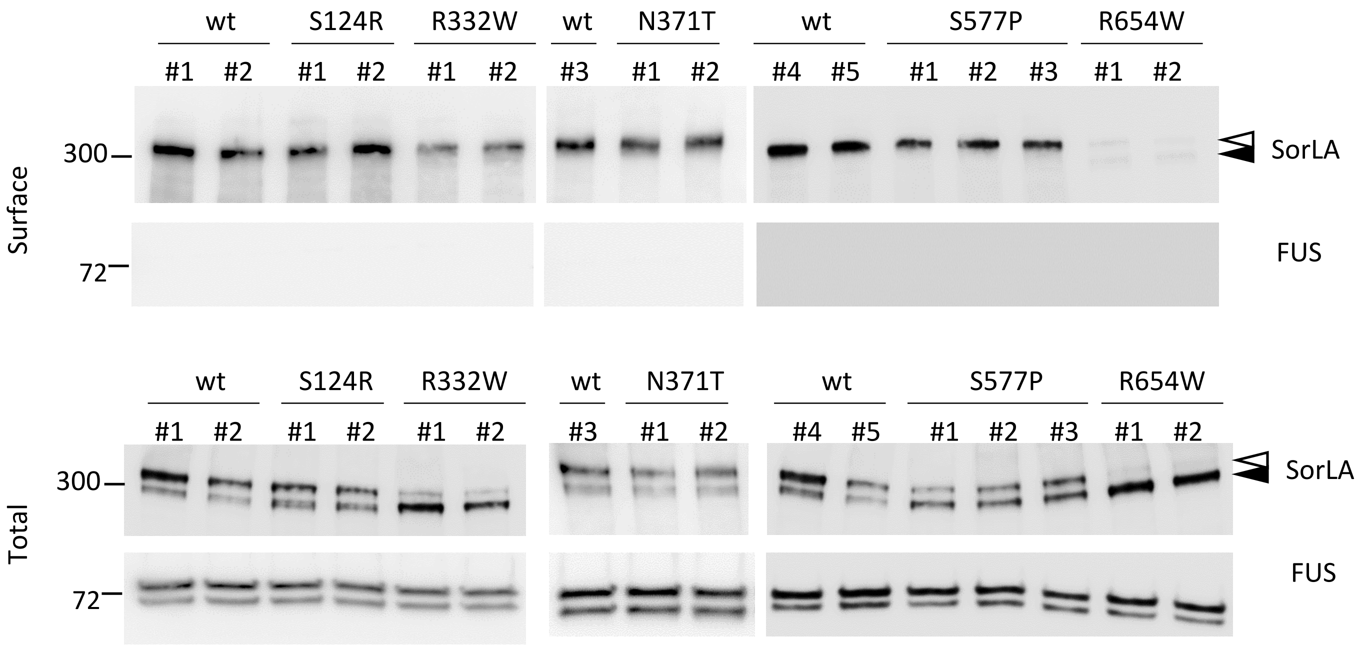

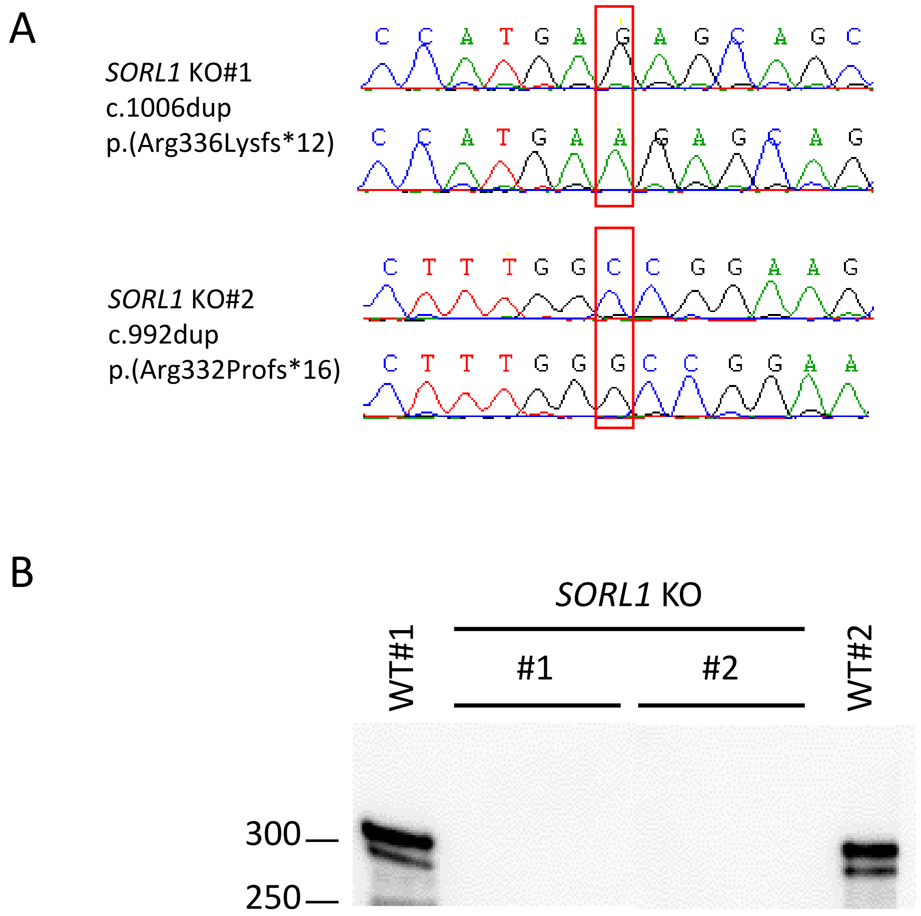

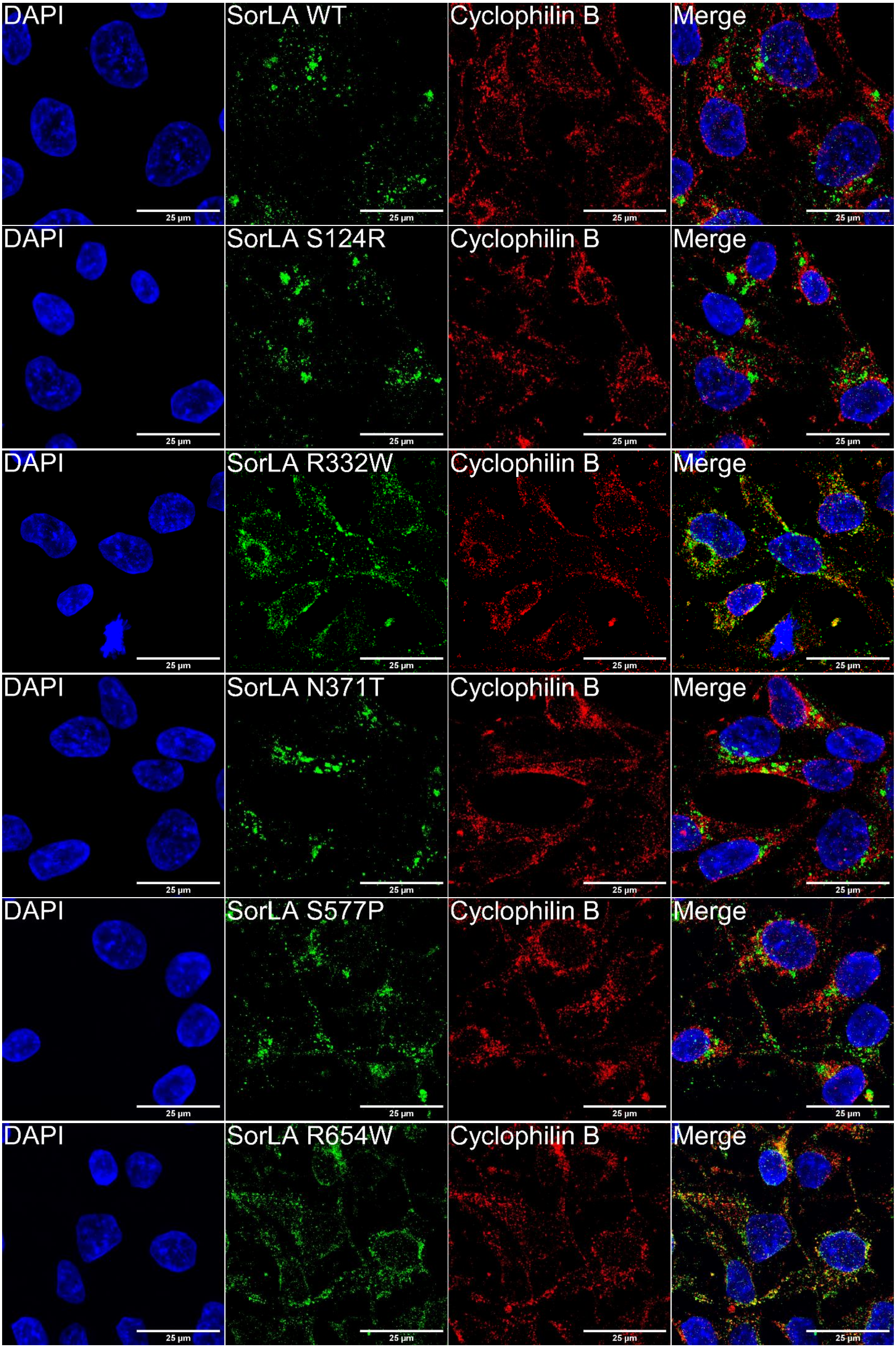

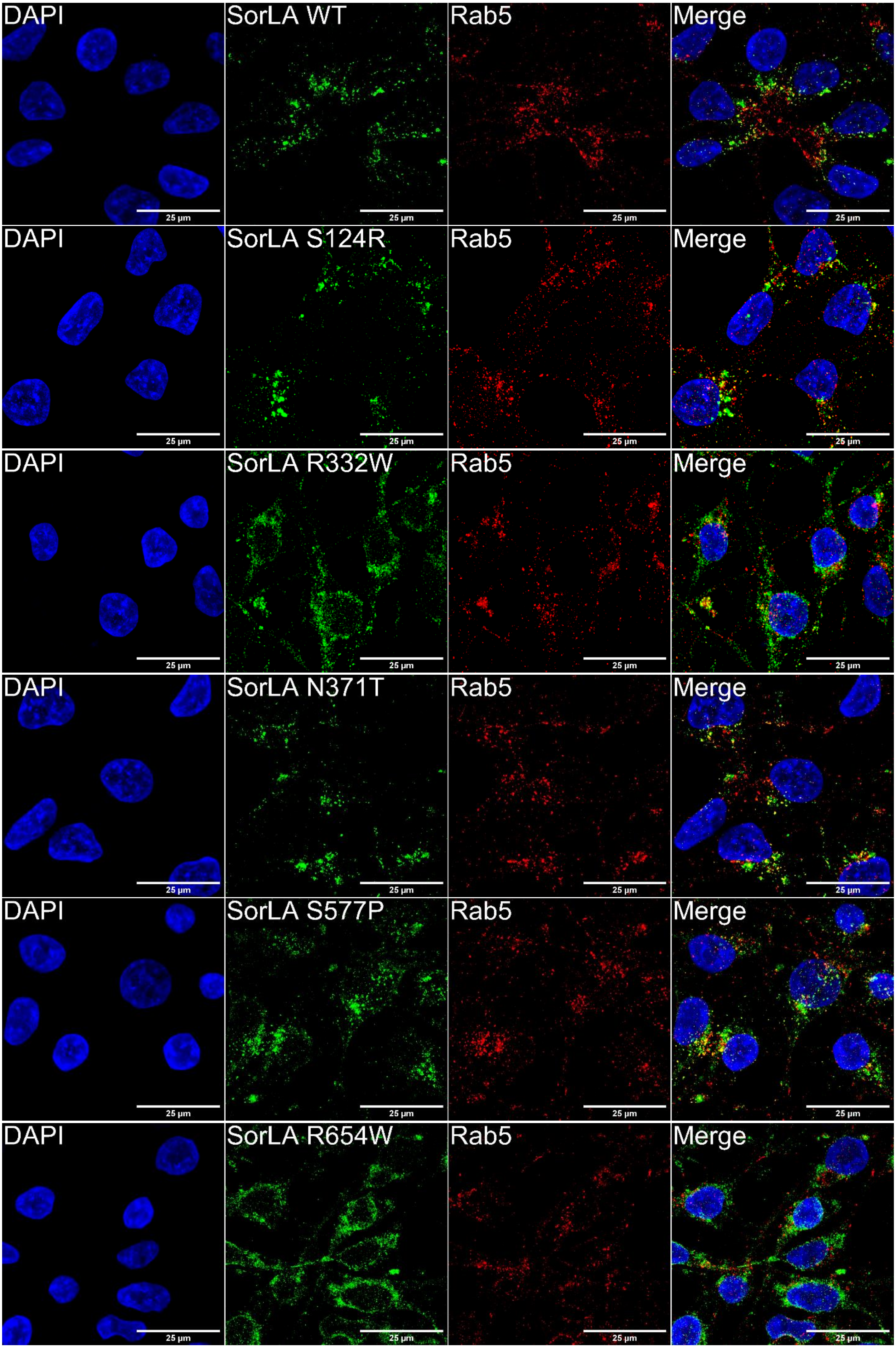

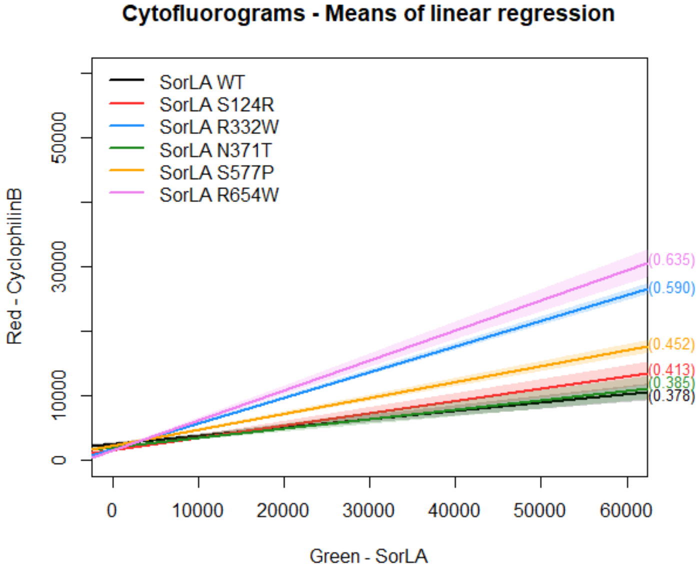

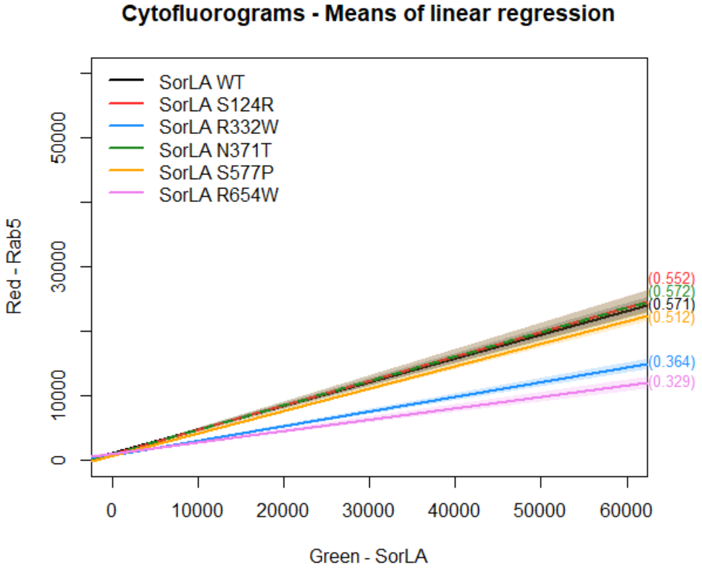
