## Supplementary material for "Impaired SorLA maturation and trafficking as a new mechanism for *SORL1* missense variants in Alzheimer disease": Table S1

| | | | HEK293 | | hiPS | | | |
|---|---|---|---|---|---|---|---|---|
| BaseChange | AAChange | Functional Domain | Maturation defect | Trafficking defect | Maturation defect | Reduced Cell surface expression | ER retention | Aβ secretion |
| c.160G>C | p.Asp54His | Propeptide | N | | | | | |
| c.201G>T | p.Arg67Ser | Propeptide | N | | | | | |
| c.227C>T | p.Pro76Leu | Propeptide | N | N | | | | |
| c.340A>C | p.Ser114Arg | LinkerPropeptideVPS10P | Y | Y | | | | |
| c.372C>G | p.Ser124Arg | LinkerPropeptideVPS10P | N | N | N | N | N | N |
| c.418G>A | p.Asp140Asn | VPS10P | N | N | | | | |
| c.422A>G | p.Tyr141Cys | VPS10P | N | N | | | | |
| c.994C>T | p.Arg332Trp | VPS10P | Y | Y | Y | Y | Y | Y |
| c.1112A>C | p.Asn371Thr | VPS10P | N | N | N | N | N | N |
| c.1418G>C | p.Cys473Ser | VPS10P | N | N | | | | |
| c.1469G>T | p.Arg490Leu | VPS10P | N | N | | | | |
| c.1531G>C | p.Gly511Arg | VPS10P | N | N | | | | |
| c.1628G>A | p.Gly543Glu | VPS10P | Y | Y | | | | |
| c.1690A>G | p.Ser564Gly | VPS10P | Y | Y | | | | |
| c.1729T>C | p.Ser577Pro | LinkerVPS10PSortilinC | Y | Y | Y | N | Y | N |
| c.1805C>T | p.Ser602Leu | SortilinC | N | N | | | | |
| c.1960C>T | p.Arg654Trp | SortilinC | Y | Y | Y | Y | Y | Y |
| c.2021A>G | p.Asn674Ser | SortilinC | N | | | | | |
| c.2185C>T | p.Arg729Trp | SortilinC | Y | Y | | | | |
| c.2311C>T | p.Arg771Cys | LinkerSortilinCBetaPropeller | N | | | | | |
| c.2362C>T | p.Arg788Trp | LinkerSortilinCBetaPropeller | N | | | | | |
| c.2416G>A | p.Asp806Asn | YWTD-Beta propeller | Y | Y | | | | |
| c.2629C>T | p.Arg877Cys | YWTD-Beta propeller | N | | | | | |
| c.2650G>A | p.Val884Met | YWTD-Beta propeller | N | | | | | |
| c.2771A>G | p.Asn924Ser | YWTD-Beta propeller | N | | | | | |
| c.2801A>G | p.Tyr934Cys | YWTD-Beta propeller | Y | Y | | | | |
| c.2956G>A | p.Ala986Thr | YWTD-Beta propeller | N | | | | | |
| c.2995A>G | p.Asn999Asp | YWTD-Beta propeller | N | | | | | |
| c.3285G>C | p.Trp1095Cys | CR-cluster Class A repeats | N | | | | | |
| c.3295T>C | p.Phe1099Leu | CR-cluster Class A repeats | N | | | | | |
| c.3346A>G | p.Ile1116Val | CR-cluster Class A repeats | N | | | | | |
| c.3436G>A | p.Asp1146Asn | CR-cluster Class A repeats | N | | | | | |
| c.3483C>G | p.Asp1161Glu | CR-cluster Class A repeats | N | | | | | |
| c.3658G>A | p.Gly1220Arg | CR-cluster Class A repeats | N | | | | | |
| c.3727C>T | p.Arg1243Cys | CR-cluster Class A repeats | N | | | | | |
| c.3763C>T | p.His1255Tyr | CR-cluster Class A repeats | N | | | | | |
| c.3827C>T | p.Thr1276Met | CR-cluster Class A repeats | N | | | | | |
| c.3892G>A | p.Gly1298Arg | CR-cluster Class A repeats | N | | | | | |
| c.3907C>T | p.Arg1303Cys | CR-cluster Class A repeats | N | | | | | |
| c.3929C>T | p.Ala1310Val | CR-cluster Class A repeats | N | | | | | |
| c.4051G>A | p.Gly1351Ser | CR-cluster Class A repeats | N | | | | | |
| c.4073A>G | p.Asn1358Ser | CR-cluster Class A repeats | N | | | | | |
| c.4166A>T | p.Asp1389Val | CR-cluster Class A repeats | N | | | | | |
| c.4176C>A | p.Asn1392Lys | CR-cluster Class A repeats | N | | | | | |
| c.4265A>G | p.Asn1422Ser | CR-cluster Class A repeats | N | | | | | |
| c.4303A>T | p.Thr1435Ser | CR-cluster Class A repeats | N | | | | | |
| c.4432T>A | p.Cys1478Ser | CR-cluster Class A repeats | N | | | | | |
| c.4603G>A | p.Asp1535Asn | CR-cluster Class A repeats | Y | Y | | | | |
| c.4629G>C | p.Glu1543Asp | CR-cluster Class A repeats | N | N | | | | |
| c.4633G>A | p.Asp1545Asn | CR-cluster Class A repeats | N | | | | | |
| c.4635T>A | p.Asp1545Glu | CR-cluster Class A repeats | Y | Y | | | | |
| c.4811A>G | p.Glu1604Gly | Fibronectin-III cluster | N | N | | | | |
| c.4924G>A | p.Val1642Met | Fibronectin-III cluster | N | | | | | |
| c.4961C>T | p.Pro1654Leu | Fibronectin-III cluster | Y | Y | | | | |
| c.4964G>A | p.Arg1655Gln | Fibronectin-III cluster | N | | | | | |
| c.5020G>A | p.Ala1674Thr | Fibronectin-III cluster | N | | | | | |
| c.5042G>A | p.Gly1681Asp | Fibronectin-III cluster | N | N | | | | |
| c.5185C>G | p.Arg1729Gly | Fibronectin-III cluster | N | N | | | | |
| c.5186G>A | p.Arg1729His | Fibronectin-III cluster | N | | | | | |
| c.5439T>A | p.His1813Gln | Fibronectin-III cluster | N | | | | | |
| c.5447A>G | p.Tyr1816Cys | Fibronectin-III cluster | Y | Y | | | | |
| c.5586G>T | p.Trp1862Cys | Fibronectin-III cluster | Y | Y | | | | |
| c.5596C>T | p.Arg1866Trp | Fibronectin-III cluster | N | | | | | |
| c.5740C>T | p.Pro1914Ser | Fibronectin-III cluster | Y | Y | | | | |
| c.5953C>T | p.Arg1985Cys | Fibronectin-III cluster | N | | | | | |
| c.6194A>T | p.Asp2065Val | Fibronectin-III cluster | N | | | | | |
| c.6224G>C | p.Gly2075Ala | Fibronectin-III cluster | N | | | | | |
| c.6248A>G | p.Lys2083Arg | Fibronectin-III cluster | N | | | | | |
| c.6289G>A | p.Val2097Ile | Fibronectin-III cluster | N | | | | | |
| c.6401C>T | p.Thr2134Met | LinkerFNIIITM | N | | | | | |
| c.6525C>A | p.Ser2175Arg | CytoplasmicTail | N | | | | | |
