## Supplementary material for "Impaired SorLA maturation and trafficking as a new mechanism for *SORL1* missense variants in Alzheimer disease": Table S2

| Variants | ΔΔG (kcal/mol) | | ΔΔG (kcal/mol) | | $\Delta\Delta S_{vib}$ (kcal/mol) | |
|---|---|---|---|---|---|---|
| | **DynaMut** | | **ENCoM** | | **ENCoM** | |
| S114R | 0,326 | Stabilizing | 0,253 | Stabilizing | -0,316 | Decreased molecular flexibility |
| S124R | 0,969 | Stabilizing | 0,648 | Stabilizing | -0,81 | Decreased molecular flexibility |
| D140N | -0,622 | Destabilizing | -0,219 | Destabilizing | 0,273 | Increased molecular flexibility |
| Y141C | -1,108 | Destabilizing | -1,055 | Destabilizing | 1,318 | Increased molecular flexibility |
| E270K | 0,251 | Stabilizing | -0,039 | Destabilizing | 0,049 | Increased molecular flexibility |
| R332W | 0,994 | Stabilizing | 0,438 | Stabilizing | -0,547 | Decreased molecular flexibility |
| N371T | -0,509 | Destabilizing | -0,098 | Destabilizing | 0,123 | Increased molecular flexibility |
| C473S | 0,618 | Stabilizing | -0,045 | Destabilizing | 0,057 | Increased molecular flexibility |
| R490L | -0,076 | Destabilizing | -0,566 | Destabilizing | 0,707 | Increased molecular flexibility |
| G511R | 0,136 | Stabilizing | 0,244 | Stabilizing | -0,305 | Decreased molecular flexibility |
| G543E | -0,484 | Destabilizing | 0,756 | Destabilizing | -0,945 | Decreased molecular flexibility |
| S564G | -0,588 | Destabilizing | -0,631 | Destabilizing | 0,789 | Increased molecular flexibility |
| S577P | -0,411 | Destabilizing | -0,183 | Destabilizing | 0,229 | Increased molecular flexibility |
| S602L | 0,805 | Stabilizing | 0,116 | Stabilizing | -0,144 | Decreased molecular flexibility |
| R654W | 0,401 | Stabilizing | 0,505 | Stabilizing | -0,631 | Decreased molecular flexibility |
| N674S | -0,271 | Destabilizing | -0,047 | Destabilizing | 0,059 | Increased molecular flexibility |
| R729W | -0,083 | Destabilizing | -0,077 | Destabilizing | 0,096 | Increased molecular flexibility |
