## Supplementary material for "Impaired SorLA maturation and trafficking as a new mechanism for *SORL1* missense variants in Alzheimer disease": Table S3

| BaseChange | AAChange | AAChange | | Oligonucleotide Sequence (5'...3') |
|---|---|---|---|---|
| c.160G>C | p.Asp54His | D54H | 160 F | TCGTGGTGCAGGGC**C**ACCCGCGCGAGCT |
| | | | 160 R | AGCTCGCGCGGGT**G**GCCCTGCACCACGA |
| c.201G>T | p.Arg67Ser | R67S | 201F | CGGGGATGCCAG**C**GGGGCGAGCC |
| | | | 201R | GGCTCGCCCCGCTGGCATCCCCG |
| c.227C>T | p.Pro76Leu | P76L | 227_F | GCGGACGAGAAGC**T**GCTCCGGAGGAAACG |
| | | | 227_R | CGTTTCCTCCGGAGC**A**GCTTCTCGTCCGC |
| c.340A>C | p.Ser114Arg | S114R | 340_F | GGCTGGAGAGAAA**C**GCAACGTGATCGTGG |
| | | | 340_R | CCACGATCACGTTGC**G**TTTCTCTCCAGCC |
| c.372C>G | p.Ser124Arg | S124R | 372 F | CTTGGCCCGAGATAG**G**CTGGCATTGGCGAGG |
| | | | 372 R | CCTCGCCAATGCCAG**C**CTATCTCGGGCCAAG |
| c.418G>A | p.Asp140Asn | D140N | 418 F | GCAGTGATGTGTACGTGTCTTAC**A**ACTATGGAAAATCATTCAAG |
| | | | 418 R | CTTGAATGATTTTCCATAGT**T**GTAAGACACGTACACATCACTGC |
| c.422A>G | p.Tyr141Cys | Y141C | 422 F | GATGTGTACGTGTCTTACGACT**G**TGGAAAATCATTCAAG |
| | | | 422 R | CTTGAATGATTTTCCA**C**AGTCGTAAGACACGTACACATC |
| c.808G>A | p.Glu270Lys | E270K | 808F | TCTACATTGAACGACAT**A**AACCCTCTGGCTACTCC |
| | | | 808R | GGAGTAGCCAGAGGGTTTATGTCGTTCAATGTAGA |
| c.994C>T | p.Arg332Trp | R332W | 994 F | GGTCTCCTTTGGC**T**GGAAGCCCATGAGAG |
| | | | 994 R | CTCTCATGGGCTTCC**A**GCCAAAGGAGACC |
| c.1112A>C | p.Asn371Thr | N371T | 1112_F | CACAGTAACAACCGCACCA**C**TTTATACATCTCAGAGG |
| | | | 1112_R | CCTCTGAGATGTATAAA**G**TGGTGCGGTTGTTACTGTG |
| c.1418G>C | p.Cys473Ser | C473S | 1418F | AGCTTTCCCAGGGCT**C**TTCCCTTCATCTGGC |
| | | | 1418R | GCCAGATGAAGGGAAGAGCCCTGGGAAAGCT |
| c.1469G>T | p.Arg490Leu | R490L | 1469_F | CCTCAACCTCCAGCTCC**T**GAGAATGCCCATCC |
| | | | 1469_R | GGATGGGCATTCTC**A**GGAGCTGGAGGTTGAGG |
| c.1531G>C | p.Gly511Arg | G511R | 1531 F | CCACTGGCTCAGTG**C**GAAAGAACTTGGCTAGC |
| | | | 1531 R | GCTAGCCAAGTTCTTTC**G**CACTGAGCCAGTGG |
| c.1628G>A | p.Gly543Glu | G543E | 1628F | CACTACTACACATGGG**A**AGACCACGGCGGAATC |
| | | | 1628R | GATTCCGCCGTGGTCT**T**CCCATGTGTAGTAGTG |
| c.1690A>G | p.Ser564Gly | S564G | 1690_F | CCAACGAGCTAAAATAC**G**GTACCAATGAAGGGGAG |
| | | | 1690_R | CTCCCCTTCATTGGTAC**C**GTATTTTAGCTCGTTGG |
| c.1729T>C | p.Ser577Pro | S577P | 1729_F | GGAAAACATTCATCTTC**C**CTGAGAAGCCAGTGTTTGTG |
| | | | 1729_R | CACAAACACTGGCTTCTCAG**G**GAAGATGAATGTTTTCC |
| c.1805C>T | p.Ser602Leu | S602L | 1805_F | GTCTTCACCATCTTTGGCT**T**GAACAAAGAGAATGTCC |
| | | | 1805_R | GGACATTCTCTTTGTTC**A**AGCCAAAGATGGTGAAGAC |
| c.1960C>T | p.Arg654Trp | R654W | 1960 F | GACTGTTTTCAAACGG**T**GGACCCCCCATGCCACATG |
| | | | 1960 R | CATGTGGCATGGGGGGTCC**A**CCGTTTGAAAACAGTC |
| c.2021A>G | p.Asn674Ser | N674S | 2021F | GGTGGTCGTGTCCA**G**CTGCTCCTGCACC |
| | | | 2021R | GGTGCAGGAGCAG**C**TGGACACGACCACC |
| c.2185C>T | p.Arg729Trp | 729 | 2185 F | GAGAACGAGAGGCTAC**T**GGAAGATTTCTGGGGAC |
| | | | 2185 R | GTCCCCAGAAATCTTCC**A**GTAGCCTCTCGTTCTC |
| c.2311C>T | p.Arg771Cys | R771C | 2311_F | GTGAGGAAATCCATCTAC**T**GCTATGACCTGGCC |
| | | | 2311_R | GGCCAGGTCATAGC**A**GTAGATGGATTTCCTCAC |
| c.2362C>T | p.Arg788Trp | R788W | 2362_F | CCTCTCACCGGGCTA**T**GGGCAGCAGTGGC |
| | | | 2362_R | GCCACTGCTGCCC**A**TAGCCCGGTGAGAGG |
| c.2416G>A | p.Asp806Asn | D806N | 2416F | ACTGTTTGTATTGGTCC**A**ACCTGGCCTTGGACGTC |
| | | | 2416R | GACGTCCAAGGCCAGGTTGGACCAATACAAACAGT |
| c.2629C>T | p.Arg877Cys | R877C | 2629_F | CCTCTGTGCTTGAT**T**GTCCCAGGGCTCTGG |
| | | | 2629_R | CCAGAGCCCTGGGAC**A**ATCAAGCACAGAGG |
| c.2650G>A | p.Val884Met | V884M | 2650 F | GGGCTCTGGTCCTC**A**TGCCCCAAGAGGGGGTG |
| | | | 2650 R | CACCCCCTCTTGGGGCA**T**GAGGACCAGAGCCC |
| c.2771A>G | p.Asn924Ser | 924 | 2771 F | GATGTGAAGTGGCCCA**G**TGGCATCTCTGTGGAC |
| | | | 2771 R | GTCCACAGAGATGCCA**C**TGGGCCACTTCACATC |
| c.2801A>G | p.Tyr934Cys | Y934C | 2801_F | GGACGACCAGTGGATTT**G**CTGGACGGATGCC |
| | | | 2801_R | GGCATCCGTCCAG**C**AAATCCACTGGTCGTCC |
| c.2956G>A | p.Ala986Thr | A986T | 2956_F | CAGCTCAGCATATTCCGA**A**CTTCCAAATACAGTGG |
| | | | 2956_R | CCACTGTATTTGGAAG**T**TCGGAATATGCTGAGCTG |
| c.2995A>G | p.Asn999Asp | N999D | 2995F | AGATGGAGATTCTGGCA**G**ACCAGCTCACGGG |
| | | | 2995R | CCCGTGAGCTGGTCTGCCAGAATCTCCATCT |
| c.3285G>C | p.Trp1095Cys | R1095C | 3285_F | CTGTATCAACAGCATTTG**C**TGGTGTGACTTTGACAACG |
| | | | 3285_R | CGTTGTCAAAGTCACACCA**G**CAAATGCTGTTGATACAG |
| c.3295T>C | p.Phe1099Leu | F1099L | 3295F | AGCATTTGGTGGTGTGAC**C**TTGACAACGACTGTGGAG |
| | | | 3295R | CTCCACAGTCGTTGTCAAGGTCACACCACCAAATGCT |
| c.3346A>G | p.Ile1116Val | D1161E | 3346_F | AACTGCCCTACCACC**G**TCTGTGACCTGGAC |
| | | | 3346_R | GTCCAGGTCACAGACGGTGGTAGGGCAGTT |
| c.3436G>A | p.Asp1146Asn | D1146N | 3436_F | CCTTGAGGATGACTGTGGA**A**ACAACAGTGATGAAAGTC |
| | | | 3436_R | GACTTTCATCACTGTTGT**T**TCCACAGTCATCCTCAAGG |
| c.3483C>G | p.Asp1161Glu | D1161E | 3483 F | CCAGTGCCGGAGTGA**G**GAGTACAACTGCAG |
| | | | 3483 R | CTGCAGTTGTACTC**C**TCACTCCGGCACTGG |
| c.3658G>A | p.Gly1220Arg | G1220R | 3658 F | GGTGGGCGTGTGAC**A**GGGATACGGACTGC |
| | | | 3658 R | GCAGTCCGTATCCC**T**GTCACACGCCCACC |
| c.3727C>T | p.Arg1243Cys | R1243C | 3727 F | GAAGTGCAATGGATTC**T**GCTGCCCAAACGGCAC |
| | | | 3727 R | GTGCCGTTTGGGCAGC**A**GAATCCATTGCACTTC |

| BaseChange | AAChange | AAChange | | Oligonucleotide Sequence (5'...3') |
|---|---|---|---|---|
| c.3763C>T | p.His1255Tyr | H1255Y | 3763F | TGCATCCCATCCAGCAAA**T**ATTGTGATGGTCTGCGTG |
| | | | 3763R | CACGCAGACCATCACAAT**A**TTTGCTGGATGGGATGCA |
| c.3827C>T | p.Thr1276Met | T1276M | 3827_F | GCGAGCCCCTCTGTA**T**GCACTTCATGGAC |
| | | | 3827_R | GTCCATGAAGTGC**A**TACAGAGGGGCTCGC |
| c.3892G>A | p.Gly1298Arg | G1298R | 3892 F | CTCCATGGTCTGTGAC**A**GAATCATCCAGTGCCG |
| | | | 3892R | CGGCACTGGATGATTC**T**GTCACAGACCATGGAG |
| c.3907C>T | p.Arg1303Cys | R1303C | 3907_F | GGAATCATCCAGTGC**T**GCGACGGGTCCGATG |
| | | | 3907_R | CATCGGACCCGTCGC**A**GCACTGGATGATTCC |
| c.3929C>T | p.Ala1310Val | A1310V | 3929 F | GGGTCCGATGAGGATG**T**GGCGTTTGCAGGATGC |
| | | | 3929 R | GCATCCTGCAAACGCC**A**CATCCTCATCGGACCC |
| c.4051G>A | p.Gly1351Ser | G1351S | 4051_F | GGGATGGATGATTGC**A**GCGATTATTCTGATGAAGC |
| | | | 4051_R | GCTTCATCAGAATAATCGC**T**GCAATCATCCATCCC |
| c.4073A>G | p.Asn1358Ser | N1358S | 4073 F | CTGATGAAGCCA**G**CTGCGAAAACCCCACAG |
| | | | 4073 R | CTGTGGGGTTTTCGCAG**C**TGGCTTCATCAG |
| c.4166A>T | p.Asp1389Val | D1389V | 4166 F | CAACAGATGGAAATGTG**T**CAGGGAGAACGACTGTGG |
| | | | 4166 R | CCACAGTCGTTCTCCCTG**A**CACATTTCCATCTGTTG |
| c.4176C>A | p.Asn1392Lys | N1392K | 4176_F | GGAAATGTGACAGGGAGAA**G**GACTGTGGGGACTGG |
| | | | 4176_R | CCAGTCCCCACAGTC**C**TTCTCCCTGTCACATTTCC |
| c.4265A>G | p.Asn1422Ser | N1422S | 4265F | ACGTGTCTGCCCA**G**TTACTACCGCTGCAG |
| | | | 4265R | CTGCAGCGGTAGTAACTGGGCAGACACGT |
| c.4303A>T | p.Thr1435Ser | T1435S | 4303 F | ACCTGCGTGATGGAC**T**CCTGGGTGTGCGACG |
| | | | 4303 R | CGTCGCACACCCAGG**A**GTCCATCACGCAGGT |
| c.4432T>A | p.Cys1478Ser | C1478S | 4432_F | CGATTTGAGTTCGAA**A**GCCACCAACCGAAGACG |
| | | | 4432_R | CGTCTTCGGTTGGTGGC**T**TTCGAACTCAAATCG |
| c.4603G>A | p.Asp1535Asn | D1535N | 4603_F | CTCGGAGCGCTGC**A**ACGGCTTCCTGGAC |
| | | | 4603_R | GTCCAGGAAGCCGT**T**GCAGCGCTCCGAG |
| c.4629G>C | p.Glu1543Asp | E1543D | 4629F | TGGACTGCTCGGACGA**C**AGCGATGAAAAGGCCTG |
| | | | 4629R | CAGGCCTTTTCATCGCTGTCGTCCGAGCAGTCCA |
| c.4635T>A | p.Asp1545Glu | D1545E | 4635_F | GCTCGGACGAGAGCGA**A**GAAAAGGCCTGCAGTG |
| | | | 4635_R | CACTGCAGGCCTTTTC**T**TCGCTCTCGTCCGAGC |
| c.4633G>A | p.Asp1545Asn | D1545N | 4633_F | CTGCTCGGACGAGAGC**A**ATGAAAAGGCCTGCAG |
| | | | 4633_R | CTGCAGGCCTTTTCAT**T**GCTCTCGTCCGAGCAG |
| c.4811A>G | p.Glu1604Gly | E1604G | 4811 F | ATATGGAAGACTCTGG**G**GACCCACAGCAATAAG |
| | | | 4811 R | CTTATTGCTGTGGGTC**C**CCAGAGTCTTCCATAT |
| c.4924G>A | p.Val1642Met | V1642M | 4924 F | CAACACCAATGACTTT**A**TGACCCTGAGGACCCC |
| | | | 4924 R | GGGGTCCTCAGGGTCA**T**AAAGTCATTGGTGTTG |
| c.4961C>T | p.Pro1654Leu | P1654L | 4961 F | GATTGCCAGATGCCC**T**TCGAAATCTCCAGCTGTC |
| | | | 4961 R | GACAGCTGGAGATTTCGA**A**GGGCATCTGGCAATC |
| c.4964G>A | p.Arg1655Gln | R1655Q | 4964F | GATTGCCAGATGCCCCTC**A**AAATCTCCAGCTGTCA |
| | | | 4694R | TGACAGCTGGAGATTTTGAGGGGCATCTGGCAATC |
| c.5020G>A | p.Ala1674Thr | A1674T | 5020F | TGATTGTAGGCCACTGG**A**CTCCTCCCATCCACA |
| | | | 5020R | TGTGGATGGGAGGAGTCCAGTGGCCTACAATCA |
| c.5042G>A | p.Gly1681Asp | G1681D | 5042 F | CCATCCACACCCATG**A**CCTCATCCGTGAGTAC |
| | | | 5042 R | GTACTCACGGATGAGG**T**CATGGGTGTGGATGG |
| c.5186G>A | p.Arg1729His | R1729H | 5186 F | GCTGCGGTGACTAGTC**A**TGGAATAGGAAACTGG |
| | | | 5186 R | CCAGTTTCCTATTCCA**T**GACTAGTCACCGCAGC |
| c.5185C>G | p.Arg1729Gly | R1729G | 5185F | TGGCTGCGGTGACTAGT**G**GTGGAATAGGAAACTGG |
| | | | 5185R | CCAGTTTCCTATTCCACCACTAGTCACCGCAGCCA |
| c.5439T>A | p.His1813Gln | H1813Q | 5439 F | GTTGGCAATCTGACAGCTCA**A**ACATCCTATGAGATTTCTG |
| | | | 5439 R | CAGAAATCTCATAGGATGT**T**TGAGCTGTCAGATTGCCAAC |
| c.5447A>G | p.Tyr1816Cys | Y1816C | 5447_F | GACAGCTCATACATCCT**G**TGAGATTTCTGCCTGG |
| | | | 5447_R | CCAGGCAGAAATCTCA**C**AGGATGTATGAGCTGTC |
| c.5586G>T | p.Trp1862Cys | W1862C | 5586_F | GCAGTGGAATGTACCTG**T**ACCGGCCCCCGGAATG |
| | | | 5586_R | CATTCCGGGGGCCGGT**A**CAGGTACATTCCACTGC |
| c.5596C>T | p.Arg1866Trp | R1866W | 5596 F | TACCTGGACCGGCCCC**T**GGAATGTGGTTTATGG |
| | | | 5596 R | CCATAAACCACATTCC**A**GGGGCCGGTCCAGGTA |
| c.5740C>T | p.Pro1914Ser | P1914S | 5740_F | GTCCGTGTAGTGGTA**T**CCTACCAGGGGCCATC |
| | | | 5740_R | GATGGCCCCTGGTAGG**A**TACCACTACACGGAC |
| c.5953C>T | p.Arg1985Cys | R1985C | 5953F | GGAGCTACAAAGTAAAATCC**T**GTAACAGCACTGTGGAATAC |
| | | | 5953R | GTATTCCACAGTGCTGTTACAGGATTTTACTTTGTAGCTCC |
| c.6194A>T | p.Asp2065Val | D2065V | 6194 F | GGCTATGAGATACACATGTTTG**T**TAGTGCCATGAATATCACAGC |
| | | | 6194 R | GCTGTGATATTCATGGCACTA**A**CAAACATGTGTATCTCATAGCC |
| c.6224G>C | p.Gly2075Ala | G2075A | 6224 F | CCATGAATATCACAGCTTACCTTG**C**GAATACTACTGACAATTTC |
| | | | 6224 R | GAAATTGTCAGTAGTATTC**G**CAAGGTAAGCTGTGATATTCATGG |
| c.6248A>G | p.Lys2083Arg | K2083R | 6248 F | CTACTGACAATTTCTTTA**G**AATTTCCAACCTGAAGATGGGTC |
| | | | 6248 R | GACCCATCTTCAGGTTGGAAATT**C**TAAAGAAATTGTCAGTAG |
| c.6289G>A | p.Val2097Ile | V2097I | 6289 F | AATTACACGTTCACC**A**TCCAAGCAAGATGCC |
| | | | 6289 R | GGCATCTTGCTTGGA**T**GGTGAACGTGTAATT |
| c.6401C>T | p.Thr2134Met | T2134M | 6401 F | GCAGGCTGCCAGATCTA**T**GGATGTTGCTGCTG |
| | | | 6401 R | CAGCAGCAACATCC**A**TAGATCTGGCAGCCTGC |
| c.6525C>A | p.Ser2175Arg | S2175R | 6525_F | GCCTTCGCCAACAG**A**CACTACAGCTCCAGG |
| | | | 6525_R | CCTGGAGCTGTAGTG**T**CTGTTGGCGAAGGC |
