## Supplementary material for "Impaired SorLA maturation and trafficking as a new mechanism for *SORL1* missense variants in Alzheimer disease": Table S4

**Chr11(GRCh37):g.121340802C>G**
**SORL1 (NM_003105):c.372C>G; p.S124R**

| | |
|---|---|
| gRNA35-F | **TGCCAGGCTATCTCGGGCCA** |
| gRNA35-R | **TGGCCCGAGATAGCCTGGCA** |
| ssODN-gRNA35-antisense | ggtgtgcccagagccaacacaacacttacATCACTGCTCTTGGGCCTCGCCAATGCCAG**C**CTATCTCGGGCCAAAGCCACGATCACGTTGCTTTTCTCTCCAGCCCAGTGCACCACCATCTGATTGTGGG |
| gRNA51-F | **TCTTGGGCCTCGCCAATGCC** |
| gRNA51-R | **GGCATTGGCGAGGCCCAAGA** |
| ssODN-gRNA51-sense | GATGGTGGTGCACTGGGCTGGAGAGAAAAGCAACGTGATCGTGGCCTTGGCCCGAGATAG**G**CTGGCATTGGCGAGGCCCAAGAGCAGTGATgtaagtgttgtgttggctctgggcacacc |
| Sequencing primers | |
| Exon2-F | CTTGGCCCAAATGCTTTAGA |
| Exon2-R | TCCCTGTCATACAATTCATCCA |

**Chr11(GRCh37):g.121383766C>T**
**SORL1 (NM_003105):c.994C>T; p.R332W**

| | |
|---|---|
| gRNA53-F | **TCTCATGGGCTTCCGGCCAA** |
| gRNA53-R | **TTGGCCGGAAGCCCATGAGA** |
| ssODN-gRNA53-antisense | AGACCCCACTCACATTAATAGGATGTCTTGTGACAAACTGGGCTGCTCTCATGGGCTTCC**A**GCCAAAAGAGACCCAGAGCTGGACAGAAGACTGCTGTTCACTGCCCAAGAGATGCTGCCA |
| gRNA67-F | **ACAAACTGGGCTGCTCTCAT** |
| gRNA67-R | **ATGAGAGCAGCCCAGTTTGT** |
| ssODN-gRNA67-sense | TGGCAGCATCTCTTGGGCAGTGAACAGCAGTCTTCTGTCCAGCTCTGGGTCTCCTTTGGC**T**GGAAGCCCATGAGAGCAGCCCAGTTTGTCACAAGACATCCTATTAATGTGAGTGGGGTCT |
| Sequencing primers | |
| Exon7-F | TGCAATCACAGTCTCTATTTTGTG |
| Exon7-R | GAACCACAGACCACGATGC |

**Chr11(GRCh37):g.121384931A>C**
**SORL1(NM_003105):c.1112A>C; p.N371T**

| | |
|---|---|
| gRNA43-F | **AATTGGTGCGGTTGTTACTG** |
| gRNA43-R | **CAGTAACAACCGCACCAATT** |
| ssODN-gRNA43-antisense | CGTTCTCCAAGGACAGGGAGAACTTCAGCCCCTCTGCCTCTGAGATGTATAAA**G**TGGTGCGGTTGTTACTGTGACTGACACACACAAACACCTGGTCCTCGGAGGCATCTGCGATGTAATATTC |
| gRNA51-F | **CACCAATTTATACATCTCAG** |
| gRNA51-R | AAAC**CTGAGATGTATAAATTGGTG** |
| ssODN-gRNA43-sense | GCCTCCGAGGACCAGGTGTTTGTGTGTGTCAGCCACAGTAACAACCGCACCA**C**TTTATACATCTCAGAAGCAGAGGGGCTGAAGTTCTCCCTGTCCTTGGAGAACGTGCTCTATTACAGCCC |
| Sequencing primers | |
| Exon8-F | CAAAGCTTCTTTGCCAAGGT |
| Exon8-R | GCTCGGACCCTAAATGTTACC |

**Chr11(GRCh37):g.121414300T>C**
**SORL1 (NM_003105):c.1729T>C; p.S577P**

| | |
|---|---|
| gRNA35-F | **GTACCAATGAAGGGGAGACC** |
| gRNA35-R | **GGTCTCCCCTTCATTGGTAC** |
| ssODN-gRNA35-antisense | CTTCTCCCCAGGTTCTGTGAGGAGGCCATACACAAACACTGGCTTCTCAG**G**GAAGATGAATGTTTTCCATGTCTCTCCTTCATTGGTACTGTATctgcccaaatgcaagtaagagaatc |
| gRNA26-F | **GCAGATACAGTACCAATGAA** |
| gRNA26-R | **TTCATTGGTACTGTATCTGC** |
| ssODN-gRNA26-antisense | CTCTTCTCCCCAGGTTCTGTGAGGAGGCCATACACAAACACTGGCTTCTCAG**G**GAAGATGAATGTTTTCCAGGTCTCTCCTTCATTGGTACTGTATctgcccaaatgcaagtaagagaatccttgcagtg |
| Sequencing primers | |
| Exon13-F | AAACTTTCCCTGCCTTAGCC |
| Exon13-R | AGAATTGAGCAGGCCTTCAA |

**Chr11(GRCh37):chr11:121416047C>T**
**SORL1 (NM_003105):c.1960C>T; p.R654W**

| | |
|---|---|
| gRNA69-F | **CTCCATTGAAGCATGTGGCA** |
| gRNA69-R | **TGCCACATGCTTCAATGGAG** |
| ssODN-gRNA69-sense | TCACCATCTGATGAGCGGGGGAATGAGTGTTTGCTGGGACACAAGACTGTTTTCAAACGG**T**GGACCCCTCATGCCACATGCTTCAATGGAGAGGACTTTGACAGGCCGGTGGTCGTGTCCA |
| gRNA74-F | **GTCCTCTCCATTGAAGCATG** |
| gRNA74-R | **CATGCTTCAATGGAGAGGAC** |
| ssODN-gRNA74-sense | TCACCATCTGATGAGCGGGGGAATGAGTGTTTGCTGGGACACAAGACTGTTTTCAAACGG**T**GGACCCCCCATGCTACATGCTTCAATGGAGAGGACTTTGACAGGCCGGTGGTCGTGTCCA |
| Sequencing primers | |
| Exon14-F | AGCTGATTGGAGGGTCTGTG |
| Exon14-R | AATCTCTTCCCCCAACACTG |
