## Supplementary material for "Impaired SorLA maturation and trafficking as a new mechanism for *SORL1* missense variants in Alzheimer disease": Table S5

**P- value corresponding to Figure 3**

| | AAChange | P-value | Adjusted p-value |
|---|---|---|---|
| sSorLA$^{759}$ constructs | P76L | 4,45E-01 | 1,00E+00 |
| | S114R | 2,61E-08 | 3,65E-07 |
| | S124R | 7,82E-01 | 1,00E+00 |
| | D140N | 7,29E-01 | 1,00E+00 |
| | R332W | 5,29E-27 | 7,40E-26 |
| | C473S | 7,98E-01 | 1,00E+00 |
| | R490L | 4,38E-01 | 1,00E+00 |
| | G511R | 6,96E-01 | 1,00E+00 |
| | G543E | 1,94E-10 | 2,72E-09 |
| | S564G | 1,17E-27 | 1,64E-26 |
| | S577P | 6,94E-23 | 9,72E-22 |
| | S602L | 4,13E-01 | 1,00E+00 |
| | R654W | 3,82E-16 | 5,35E-15 |
| | R729W | 5,75E-02 | 8,05E-01 |
| sSorLA$^{2131}$ constructs | S124R | 7,50E-02 | 1,00E+00 |
| | Y141C | 6,86E-01 | 1,00E+00 |
| | R332W | 1,32E-24 | 2,24E-23 |
| | N371T | 2,53E-01 | 1,00E+00 |
| | G511R | 8,96E-02 | 1,00E+00 |
| | D806N | 1,15E-15 | 1,96E-14 |
| | Y934C | 1,41E-25 | 2,40E-24 |
| | D1535N | 1,08E-08 | 1,84E-07 |
| | E1543D | 9,17E-01 | 1,00E+00 |
| | E1545D | 6,11E-13 | 1,04E-11 |
| | E1604G | 3,17E-01 | 1,00E+00 |
| | P1654L | 2,35E-01 | 1,00E+00 |
| | G1681D | 9,83E-02 | 1,00E+00 |
| | R1729G | 4,39E-02 | 7,47E-01 |
| | Y1816C | 9,15E-02 | 1,00E+00 |
| | W1862C | 3,76E-04 | 6,39E-03 |
| | P1914S | 5,74E-05 | 9,76E-04 |

**P- value corresponding to Figure 5**

| AAChange | P-value | Adjusted p-value |
|---|---|---|
| S124R | 9,93E-01 | 1,00E+00 |
| R332W | 1,04E-04 | 5,18E-04 |
| N371T | 7,01E-01 | 1,00E+00 |
| S577P | 2,92E-02 | 1,46E-01 |
| R654W | 4,17E-12 | 2,09E-11 |

**P- value corresponding to Figure 7A**

| AAChange | P-value | Adjusted p-value |
|---|---|---|
| S124R | 1,95E-01 | 9,75E-01 |
| R332W | 1,30E-07 | 6,50E-07 |
| N371T | 7,97E-01 | 1,00E+00 |
| S577P | 3,71E-03 | 1,85E-02 |
| R654W | 3,38E-09 | 1,69E-08 |

**P- value corresponding to Figure 7B**

| AAChange | P-value | Adjusted p-value |
|---|---|---|
| S124R | 2,72E-01 | 1,00E+00 |
| R332W | 7,63E-13 | 3,82E-12 |
| N371T | 9,79E-01 | 1,00E+00 |
| S577P | 1,06E-04 | 5,31E-04 |
| R654W | 8,07E-13 | 4,04E-12 |

**P- value corresponding to Figure 8**

| AAChange | P-value | Adjusted p-value |
|---|---|---|
| KO | 3,90E-04 | 2,34E-03 |
| S124R | 6,80E-01 | 1,00E+00 |
| R332W | 2,04E-03 | 1,22E-02 |
| N371T | 4,59E-01 | 1,00E+00 |
| S577P | 3,48E-02 | 2,09E-01 |
| R654W | 1,87E-04 | 1,12E-03 |
